# Expansion of the *DFR* Gene Family in *Fagopyrum*

**DOI:** 10.1101/2025.09.20.677491

**Authors:** Syafiq Samsolnizam, Boas Pucker

## Abstract

**Background:** Gene duplication is a fundamental evolutionary mechanism that contributes to genetic innovation, such as in the diversification of plant metabolic pathways. This study investigates the duplication of dihydroflavonol-4-reductase (*DFR*) genes in *Fagopyrum*, focusing on their evolutionary origin and implications for flavonoid biosynthesis. DFR is a key enzyme in the flavonoid pathway, specifically involved in the biosynthesis of anthocyanins and proanthocyanidins. DFR can accept three main substrates: dihydrokaempferol (DHK), dihydroquercetin (DHQ), and dihydromyricetin (DHM), and they exhibit variable substrate preferences.

**Result:** Through comparative genomic analysis, this study identified three unique *DFR* gene copies in *Fagopyrum*, designated *DFR1*, *DFR2a*, and *DFR2b*. *DFR2a* and *DFR2b* show unique mutations affecting the 26-amino acid substrate-binding region, including substitution of the conserved asparagine at position 3 with valine or isoleucine, and an insertion between positions 5 and 6 involving glycine, arginine, or lysine. Synteny analysis reveals that *DFR1* is located in a conserved region across related species, while *DFR2a* and *DFR2b* lie at distinct genomic loci. Gene expression analysis shows *DFR1* is universally expressed, whereas *DFR2a* and *DFR2b* expression is primarily restricted to seeds and roots. Promoter analysis supports this, revealing the absence of MYB recognition elements in *DFR2a* and *DFR2b* that are present in the *DFR1* promoter. Selective pressure analysis indicates that *DFR2* copies are under strong purifying selection relative to ancestral *DFR1* but appear slightly relaxed between *DFR2a* and *DFR2b* in a way suggesting functional differentiation, particularly in conserved regions.

**Conclusion:** Based on multiple lines of evidence, we propose DNA duplication associated with inversion (DDAI) as a mechanism to explain the emergence of the DFR2 copies, involving a staggered single-strand break, followed by inversion and non-homologous end joining repair mechanism.

## Background

Gene duplication is one of the most important evolutionary mechanisms generating functional diversity. Theoretical studies suggest that it is far easier to evolve new functions from pre-existing ones through neofunctionalization than to develop them de novo [1]. After duplication, each gene copy can evolve independently, accumulating mutations that can potentially lead to functional divergence and innovation [1]. Gene duplication can occur through several molecular mechanisms, each producing different outcomes in terms of genomic structure. In general, gene duplication refers to the duplication of a genomic segment, which may then be inserted at the same or a different location within the genome [2]. Gene duplication thus provides raw genetic material for the evolution of novel traits and functions [3]. This is particularly relevant in plants, where duplicated genes are highly abundant [2]. Plant genomes have undergone multiple rounds of whole-genome duplication (WGD) events over the past 200 million years of angiosperm evolution, which have significantly contributed to the complexity and adaptability of plant lineages [2]. In addition to their history of WGD, plant genomes tend to evolve more rapidly and exhibit greater structural variation than those of many other eukaryotes. This has resulted in larger genome sizes and a higher degree of genetic redundancy. The high abundance of gene duplicates in plants is especially interesting because of their frequent contribution to lineage-specific innovations [2].

The most extensive form of duplication is WGD, where the entire set of chromosomes is duplicated. Ancient WGD events have been detected in the ancestors of seed plants (approximately 340 million years ago) and angiosperms (around 170 million years ago) [4]. This type of duplication leads to the presence of multiple copies of every gene in the genome and is referred to as polyploidization. Polyploidy can result from hybridization between species (allopolyploidy) or occur within a single species (autopolyploidy) [5]. On a smaller scale, tandem duplications involve the replication of genes in close physical proximity, forming tandemly arrayed genes (TAGs). These local duplications typically arise through unequal crossing over during meiosis. The size of the duplicated region can vary depending on the site of chromosomal breakage. Such events may be caused by homologous recombination between similar sequences on homologous chromosomes or sister chromatids, or by non-homologous recombination resulting from replication-dependent chromosome breakage [6, 7]. In maize, for example, thousands of tandem duplicates have been identified, representing approximately 10% of all annotated genes [8]. These duplications contribute to genetic diversity and often play roles in gene family expansion and adaptive evolution. Transposable elements (TEs) can also drive sequence duplication through various mechanisms. TEs are repetitive DNA sequences with the ability to move themselves or even nearby genes to new locations within the genome.

This process, called transposition, can occur within the same chromosome or between different chromosomes. The proportion of TEs varies widely across organisms, from about 3% of the genome in yeast to more than 80% in maize [9, 10]. In *Fagopyrum esculentum*, TEs occupy a particularly large portion of the genome [11]. This includes both gradually accumulated elements and several classes of recently amplified TEs that underwent rapid multiplication between 0.5 and 1 million years ago. This transposon activity has significantly contributed to the threefold increase in genome size of *F. esculentum* compared to its close relative *Fagopyrum tataricum* [11]. There are two primary mechanisms by which TEs can cause gene duplication: retroduplication and transduplication. Retroduplication occurs when the mRNA of a host gene is reverse-transcribed into complementary DNA and inserted into a different genomic location by retrotransposon enzymes. These retrocopies are characterized by the absence of introns, the presence of a poly-A tail at the 3’ end, and flanking target site duplications (TSDs) [12]. Transduplication is facilitated by DNA transposons, particularly those with terminal inverted repeats (TIRs). These elements use a “cut-and-paste” mechanism to excise themselves and often adjacent gene sequences. They then insert into a new location in the genome, also flanked by TSDs. Unlike retrocopies, the duplicated fragments from DNA transposons can retain intron-exon structures [7, 13]. Helitrons, another type of DNA transposon, operate through a “peel-and-paste” mechanism. In this process, the positive DNA strand is peeled off to form a circular DNA molecule, which is then replicated through rolling circle replication. This mechanism can also mobilize gene fragments that may contain introns, contributing to structural genome variation [7, 13]. In addition to TE-driven duplications, segmental duplications represent another important form of gene duplication. Samonte and Eichler define segmental duplication as long stretches of duplicated sequences ranging from 1 to 200 kilobases and sharing more than 95% sequence identity [14]. Segmental duplications can arise from the transposition of chromosomal regions and often include multiple genes with intron-exon structures. Unlike tandem duplications, these segments are typically dispersed throughout the genome. This distribution suggests that mechanisms other than unequal crossing over, such as duplicative transposition, are responsible. Additional research is required to find a definitive explanation for the frequency, organization, and distribution of segmental duplications, particularly in large and complex genomes like the human genome [14].

After gene duplication, the newly formed duplicates can follow several evolutionary trajectories: (1) pseudogenization and loss, (2) neofunctionalization, or (3) subfunctionalisation. The most common fate is pseudogenization often followed by complete loss of one copy [15]. Typically, only one gene copy remains under purifying selection to maintain its original function, while the other is free to accumulate deleterious mutations that can cause it to be nonfunctional [7]. If two duplicate genes are functionally redundant and completely identical, there is generally no selective disadvantage in deleting either copy [2]. Pseudogenes are identified based on their sequence similarity to functional genes and the presence of disabling mutations. Interestingly, pseudogenes are not always rapidly eliminated. In species such as *Arabidopsis thaliana* and rice, thousands of pseudogenes are conserved within the genome [16]. One possible explanation is that some pseudogenes were only recently pseudogenized and are still in the early stages of degeneration [2]. Another possible fate is neofunctionalization, where the duplicate acquires a novel function. If this novel function provides a selective advantage, natural selection may act to preserve the gene. Ohno originally proposed the neofunctionalization model, in which one copy retains the ancestral function while its paralog evolves a new one [2, 3]. A notable example is the evolution of flavone synthase I (*FNS I*) in the *Apiaceae*, which arose through duplication of flavanone 3-hydroxylase (*F3H*), followed by neofunctionalization, as revealed by phylogenetic and comparative genomic analyses [17]. In contrast, subfunctionalisation occurs when both duplicates partition the functions of the original gene due to the accumulation of degenerative mutations. This process, also known as the duplication-degeneration-complementation (DDC) model [18], involves each copy losing a subset of ancestral functions. As a result, both gene copies must be retained to maintain the full spectrum of original functions. Subfunctionalisation does not require the evolution of new functions, but rather a division of pre-existing ones [2]. Another important trajectory is escape from adaptive conflict (EAC). Here, a single-copy gene is constrained by pleiotropy from optimizing both its ancestral and novel functions. After duplication, one copy can specialize in the ancestral role, while the other improves the novel function [19]. Unlike neofunctionalization, which involves adaptive change in only one copy (with the other maintained by purifying selection), EAC involves adaptive changes in both duplicates, with the ancestral function itself also being improved. EAC may also involve a combination of both neo- and sub-functionalisation mechanisms as it invokes both the neofunctionalization and DDC models to explain duplicate retention [2]. Although theoretically important, EAC remains understudied and can be difficult to distinguish from neofunctionalization. Other potential fates include gene dosage retention, where both duplicates do not diverge much, remain functional and are expressed to increase the overall dosage of gene product [3]. This can be advantageous for meeting metabolic demands. Additional mechanisms that influence duplicate gene retention include dosage balance and paralog interference, and these processes are not necessarily mutually exclusive [2].

Flavonoids are one of the most important classes of plant secondary metabolites, playing key roles in plant survival, adaptation, and interactions with the environment, as well as having nutritional and medicinal value for humans [20–27]. Flavonoids are a diverse family of aromatic molecules derived from phenylalanine and malonyl-coenzyme A and they comprise six major subgroups commonly found in higher plants: chalcones, flavones, flavonols, flavandiols, anthocyanins, and condensed tannins (also known as proanthocyanidins). Among them, anthocyanins are widely known for their roles as red, purple, or blue pigments, contributing to the vivid coloration of flowers and fruits [20]. In addition to their pigmentation roles, flavonoids function as phytoalexins and antioxidants, possessing reactive oxygen species (ROS) scavenging abilities [21]. They protect plants from various biotic and abiotic stresses, including UV irradiation, cold stress, pathogen infection, and herbivory [22–25]. Flavonoids also have widespread applications in food such as natural colourant [26]. Anthocyanins for example, are edible pigments that influence the taste and color of food and wine [20]. Flavones play important roles in signaling and defense responses against pathogens [27], while proanthocyanidins contribute to seed coat pigmentation [28]. Anthocyanin biosynthesis involves several enzymatic steps. A key branch-point enzyme in this pathway is dihydroflavonol 4-reductase (DFR, EC 1.1.1.219), which catalyses the NADPH-dependent reduction of dihydroflavonols into leucoanthocyanidins [29, 30]. These leucoanthocyanidins are subsequently converted into anthocyanidins by anthocyanidin synthase (ANS) and anthocyanin-related glutathione-S-transferase (arGST) and finally into stable anthocyanins via glycosylation by glycosyltransferases (**Figure 1**) [31]. DFR catalyses the reduction of dihydrokaempferol (DHK), dihydroquercetin (DHQ), and dihydromyricetin (DHM) into their corresponding leucoanthocyanidins, which are leucopelargonidin, leucocyanidin, and leucodelphinidin, respectively [32]. Since these dihydroflavonols differ only in the number of hydroxyl groups on the B-ring (which is not the site of enzymatic action), DFR enzymes in most plants can use all three substrates. Dihydroflavonols serve as substrates not only for DFR but also for FLS, which produces flavonols such as kaempferol, quercetin, and myricetin depending on substrate availability [33]. Based on currently available information, angiosperm FLS appears to prefer DHK, while *DFR* appears to favor DHQ and DHM [34]. Hydroxylation of dihydrokaempferol into dihydroquercetin or dihydromyricetin is catalysed by flavonoid 3’-hydroxylase (F3’H) and flavonoid 3’,5’-hydroxylase (F3’5’H), respectively [35]. Substrate specificity is a critical factor in determining the balance between flavonol and anthocyanin production. A 26-amino-acid region involved in DFR substrate specificity has been identified through multiple sequence alignments [36, 37]. In particular, position 133 relative to *Arabidopsis thaliana* DFR (3rd position in the 26-amino acid region) has been shown to influence substrate preference: asparagine (N) allows acceptance of all dihydroflavonols; leucine (L) restricts substrate usage to DHK; aspartic acid (D) permits the use of DHQ and DHM; and alanine (A) results in high affinity for DHK [34].

**Figure 1:**
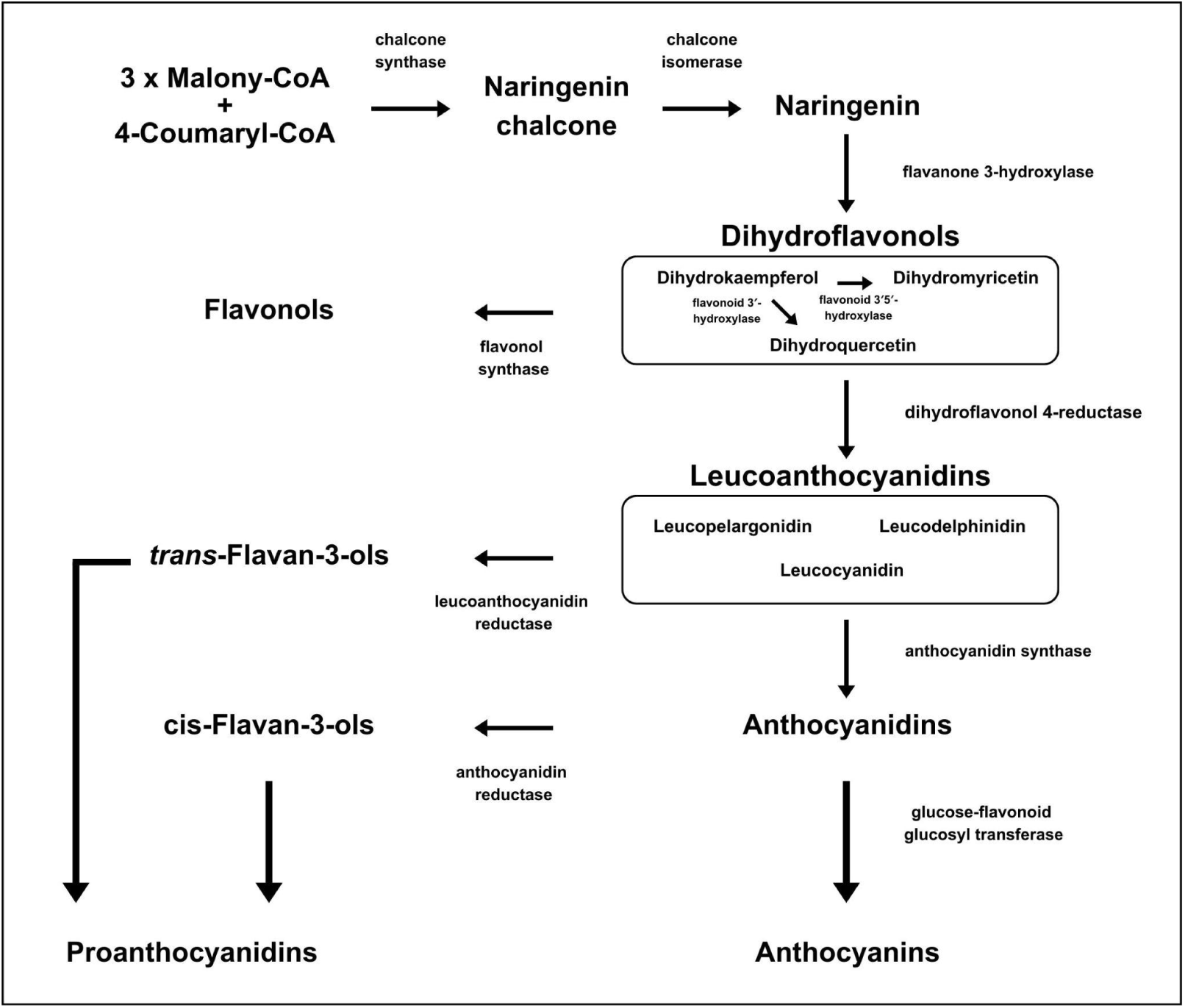
Simplified illustration of the flavonoid biosynthesis pathway. The enzyme responsible for each step is indicated next to the corresponding arrow.

The activity of different branches in flavonoid biosynthesis is tightly regulated at the transcriptional level e.g. to prevent simultaneous activity of competing branches like flavonol and anthocyanin biosynthesis [34]. In particular, *DFR* expression is activated by the MBW complex, a transcription factor complex composed of MYB, bHLH, and WD40 proteins [38].

*Fagopyrum* is a small genus comprising less than 30 species, most of which are endemic to southern China [39]. It includes two orphan crops: common buckwheat (*F. esculentum*) and tartary buckwheat (*F. tataricum*), both of which were domesticated independently from distinct wild progenitors in southwest China, outside the major centers of domestication. While *F. tataricum* is primarily cultivated in China, *F. esculentum* is grown globally, especially across Eurasia, and is valued for its rapid growth at high altitudes and its ability to thrive in low-fertility, marginal lands [40]. *Fagopyrum* is classified into two major lineages: the cymosum group (e.g., *F. esculentum*, *Fagopyrum homotropicum*, *F. tataricum*, and *Fagopyrum dibotrys* [also known as *F. cymosum*]) and the urophyllum group (e.g., *F. urophyllum*, *F. gracilipes*, and *F. leptopodum*), based on morphological characteristics and molecular phylogenetics [39, 41]. The tribe *Fagopyreae*, which contains only the genus *Fagopyrum*, is part of the family *Polygonaceae*, which is part of the order *Caryophyllales*. *Polygonaceae* consists of about 48 genera and over 1200 species, distributed worldwide but most diverse in temperate regions [42]. The plants are typically herbaceous but can also be shrubs or small trees. Within the subfamily *Polygonoideae*, major tribes include *Fagopyreae* (e.g., *Fagopyrum*), *Rumiceae* (e.g., *Rumex*, *Oxyria*), *Persicarieae* (e.g., *Persicaria*), and *Polygoneae* (e.g., *Fallopia*, *Polygonum*) [43]. *Polygonaceae* is considered a non-core family of *Caryophyllales*, a hyperdiverse order encompassing approximately 12,500 species in 39 families, accounting for about 6% of extant angiosperm diversity. The order exhibits remarkable life-history variation, from tropical trees to temperate annual herbs, from desert-adapted succulents (e.g., *Cactaceae*) to carnivorous plants [44]. This diversity makes *Caryophyllales* a compelling system for studying relationships between WGD, diversification, and adaptation to varying climates [44]. A notable feature of core *Caryophyllales* is the presence of betalain pigments that are derived from tyrosine and incorporate betalamic acid. These pigments replace the more widespread anthocyanins in some other core *Caryophyllales* lineages [45, 46]. The mutually exclusive distribution of betalains and anthocyanins has sparked decades of debate and research, revealing several mechanisms to explain this exclusivity including wholesale gene loss, modified gene function, altered gene expression, and degeneration of the MBW (MYB-bHLH-WD40) transcription factor complex regulating anthocyanin biosynthesis [47]. However, non-core *Caryophyllales* families, such as *Polygonaceae*, including the buckwheat genus *Fagopyrum*, retain the ability to synthesize anthocyanins rather than betalains [45]. Continued research into flavonoid biosynthesis within this family could provide insights into the evolutionary trajectory of pigment biosynthesis in *Caryophyllales*, and into how non-core lineages may have retained or modified ancestral anthocyanin pathways.

Katsu et al. [48] identified two copies of *DFR* genes in buckwheat using two degenerate PCR primers. These primers were designed based on conserved regions of *DFR* proteins, one of which had previously been used to isolate *DFR* homologs in other plants [49]. The two genes were named *FeDFR1a* and *FeDFR2*, where *FeDFR1a* exhibited high sequence identity (99%) to a previously deposited *F. esculentum DFR* sequence (ACZ48698), whereas *FeDFR2* showed only 80% identity [48]. Phylogenetic analysis placed *FeDFR2* in the same major clade as *FeDFR1a*, rather than with the *DFR*-like genes such as *BEN1*, suggesting that both enzymes may be involved in catalyzing the conversion of dihydroflavonols to leucoanthocyanidins . At the protein level, FeDFR1a has an asparagine residue at the third position within the substrate specificity-determining region, while FeDFR2 has an uncommon valine at the same position and contains an additional glycine residue between positions 6 and 7 (position 136 in FeDFR2) [48]. Functional assays of recombinant DFR proteins, expressed in *Escherichia coli*, were conducted to evaluate enzyme activity against three dihydroflavonol substrates and both FeDFR1a and FeDFR2 accept all three substrates; however, FeDFR1a exhibited similar catalytic efficiency for DHM and DHK, with lower efficiency for DHQ [48]. In contrast, FeDFR2 displayed approximately twice the catalytic efficiency for DHQ compared to DHM, and the lowest activity for DHK [48]. The primary goal of this study is to investigate the duplication of *DFR* genes in *F. esculentum*. By leveraging phylogenetic analysis, synteny analysis, gene expression analysis, cis-regulatory element analysis, and selective pressure analysis, this study explores the evolutionary history and mechanism of *DFR* duplication. In addition, the research examines the role of the different *DFR* genes in the anthocyanin biosynthesis pathway, which is particularly interesting in *F. esculentum* as a member of *Caryophyllales*, a lineage better known for betalain production.

## Methods

### Datasets

Genome sequences of various *Polygonaceae* species were obtained from the National Center for Biotechnology Information (NCBI). The species and their corresponding GenBank accession numbers or other sources are listed in the Additional file 1.

### Gene Prediction

Since annotations for most of the retrieved *Polygonaceae* genome sequences were not publicly available, GeMoMa v1.9 [50] was used for homology-based gene prediction. Well-annotated genome sequences and corresponding annotations from the same order of *Caryophyllales* were retrieved from the reference sequence (RefSeq) database at NCBI [51] to be used as reference in the GeMoMa pipeline. The genome sequences are *Beta vulgaris* (RefSeq: GCF_026745355.1), *Chenopodium quinoa* (RefSeq: GCF_001683475.1) and *Spinacia oleracea* (RefSeq: GCF_ 020520425.1). RNA-seq data was incorporated into the prediction to utilise intron position conservation and accurately predict the splice site for the genes.

### Alignment and Phylogenetic Tree Construction

Candidate *DFR* sequences in the *Polygonaceae* genome sequences were identified using KIPEs v3.2.6 [52] with its existing collection of bait sequences v3.3.7z. In addition, a set of *DFR* coding sequences, including *DFR*-like sequences, was retrieved from the NCBI database for phylogenetic validation. The sequences are publicly available on Github [53]. The initial candidate *DFR* sequences from *Polygonaceae* were validated through phylogenetic analysis using FastTree v2.2 [54], based on amino acid alignment generated with MAFFT v7.505 [55] using the --auto parameter. Sequences that were clustered within the main clade together with confirmed *DFR* sequences from NCBI were retained for further analysis. In contrast, sequences clustering outside of this group, such as within the *DFR*-like clade containing BAN (BANYULS/ANR) (NM_104854.4) [56], BEN1 (NM_130102.5) [57], and DRL1 (NM_119708.3) [58] from *A. thaliana*, were excluded.

The amino acid sequences and CDS from the filtered *Polygonaceae DFR*, with representatives from other *Caryophyllales* families as outgroups, were used for phylogenetic tree construction (available on Github) [53]. A codon-aware alignment approach was used for CDS sequence alignment, where the amino acid sequence was first aligned, and then substituted by their corresponding codons using a Python script [59]. The sequences were aligned using MAFFT v7.505 [55] using --maxiterate 1000 and --genafpair parameter. Maximum likelihood trees were then constructed by FastTree v2.2 [54] using standard parameters, IQ-TREE v2.4.0 [60] using the -m MFP -B 1000 parameter, and RAxML-NG v1.2.2 [61] using the --model TIM3+G4 --bs-trees 1000 --tree pars{25},rand{25} parameter. All resulting phylogenetic trees were visualised using Interactive Tree of Life (iTOL) v7 [62] and inspected for their topologies and the positions of *DFR* in *Polygonaceae* clades.

### Sequence Analysis

The retrieved Fagopyrum DFR sequences were analyzed for protein sequence similarity. A custom Python script was used to calculate sequence similarities by running multiple sequence alignment with MAFFT v7.505 [55], generating a pairwise sequence identity matrix, and visualizing the results as a heatmap [53]. The 26–amino acid conserved region was manually examined based on the MAFFT alignment and visualized using Jalview v2.11.5 [63].

### Synteny Analysis

Synteny of *DFR* in the *Polygonaceae* family was investigated using MCScan from JCVI v1.5.7 [64, 65]. Regions surrounding the *DFR* genes were manually selected for visualization in both macro- and microsynteny analyses using the JCVI toolkit. Macrosynteny visualization was performed to compare the chromosomal regions containing *DFR* in *Fagopyrum* and its relatives, using the standard parameters of MCScan. Microsynteny visualization was carried out to compare the *DFR* gene regions at the gene level, where *DFR* and its flanking genes were manually chosen for visualization. The locations of *DFR* in the macrosynteny analysis were identified manually using Jbrowse2 v3.6.4 [66]. Connections of genes between the species were manually validated and revised based on phylogenetic trees.

### Gene Expression Analysis

RNA-seq datasets were retrieved from the NCBI Sequence Read Archive (SRA) using fasterq-dump v3.0.3 [67]. Transcript quantification was performed with kallisto v0.51.1 [68] using default parameters, executed through the Python wrapper script kallisto_pipeline3.py [69]. Merged transcript TPM values and read counts were generated using merge_kallisto_output3.py [69]. Low quality or atypical datasets were removed based on TPM and read count distributions with filter_RNAseq_samples.py [69]. A custom Python script was then used to generate boxplots of *DFR* expression across tissues [53].

Because the genes of interest are highly similar and located in close proximity, RNA-seq reads were also aligned to the genome sequence with STAR v2.7.11b [70]. The results were compared with kallisto to check consistency and reduce ambiguity. Threshold parameters such as --outFilterMultimapNmax, --alignIntronMax, --outFilterScoreMin, --outFilterMatchNminOverLread, and --outFilterMismatchNoverLmax were applied. The resulting SAM files were converted into BAM format, sorted, and indexed using Samtools v1.21 [71], and manually inspected in JBrowse2 v3.6.4 [66]. Transcript quantification from the alignments was performed with the STAR --quantMode GeneCounts function.

### Promoter Region Analysis

RNA-seq reads were mapped to the genome sequence using STAR and visualized in JBrowse2 to identify the TSS. To extract the promoter regions of each *DFR* gene in *Fagopyrum*, a 1000 bp sequence upstream of the predicted transcription start site (TSS) of each gene was retrieved. The upstream regions preceding the mapped reads of each *DFR* duplicate were then extracted for further analysis. The promoter sequences were analyzed using the PLACE v30.0, PlantPan v4.0, and PlantCARE databases [72–74]. Results from the three databases were compared to identify motifs associated with *DFR* gene expression. In addition, a custom Python script was developed to visualize the output of PlantPan [53].

### Selective Pressure Analysis

The aligned coding sequences of *DFR* from *Fagopyrum* that were previously used for the alignment and phylogenetic tree construction were analysed to estimate the synonymous substitution rate (Ks) and the nonsynonymous substitution rate (Ka). The aligned sequences were first converted into an AXT file, where each sequence was compared with its orthologs or paralogs. KaKs Calculator 3.0 [75] was used to calculate the Ka/Ks ratio for each sequence pair, providing insights into the selective pressures acting on *DFR* gene duplicates. The selective pressure on duplicate gene pairs can be inferred from the Ka/Ks ratio. A Ka/Ks ratio close to 1 indicates neutral selection, which may suggest pseudogenization. A Ka/Ks ratio less than 1 (Ka/Ks < 1) indicates purifying or negative selection, meaning that the gene is being conserved. Conversely, a Ka/Ks ratio greater than 1 (Ka/Ks > 1) suggests positive selection, which may imply that the gene duplicates are evolving to acquire new functions [76]. The Ka/Ks calculations were performed using the Model Selection (MS) method [77].

### Duplication Mechanism Analysis

Several analyses were performed to investigate the mechanism behind the duplication of DFR in *Fagopyrum*. Extensive de novo TE Annotator (EDTA) v2.2.2 [78] was used on each *Fagopyrum* genome to identify TEs located near the duplicates that could have contributed to the duplication. In particular, terminal inverted repeats (TIRs) and helitrons flanking the duplicates were examined [13]. To test for segmental duplication, dot plots were generated with ModDotPlot v0.9.0 [79]. Both *DFR1* and *DFR2* loci were compared against each other and against themselves. Evidence for segmental duplication would appear as dense linear arrays of dots, either parallel or inverted relative to the main diagonal, but displaced from it [80].

## Results

### Three *DFRs are* specific to the genus of *Fagopyrum*

#### Gene Position and Structures

A total of 14 *DFR* gene copies have been identified across four *Fagopyrum* species, with three copies each in *F. esculentum*, *F. homotropicum*, and *F. dibotrys*, and 5 copies in *F. tataricum*. The three *DFR* copies in each *Fagopyrum* species show a conserved pattern and syntenic arrangement within the same chromosome (**Figure 2**). Notably, these patterns of duplicates were not observed in other species of the *Polygonaceae* family, suggesting that this gene duplication is specific to the genus of *Fagopyrum*. Additionally, *F. tataricum* possesses two extra *DFR* copies, which are located at the same genomic locus as the other *DFR* copies and remain on the same chromosome, but differ in their sequence and are assumed to be pseudogenes. *FtDFR2a* and *FtDFR2b* are located at another locus but are assumed to be caused by a genome inversion. This finding differs from the previous study by Katsu et al. [48], which identified only two *DFR* copies in *F. esculentum*, named *FeDFR1a* and *FeDFR2*.

**Figure 2:**
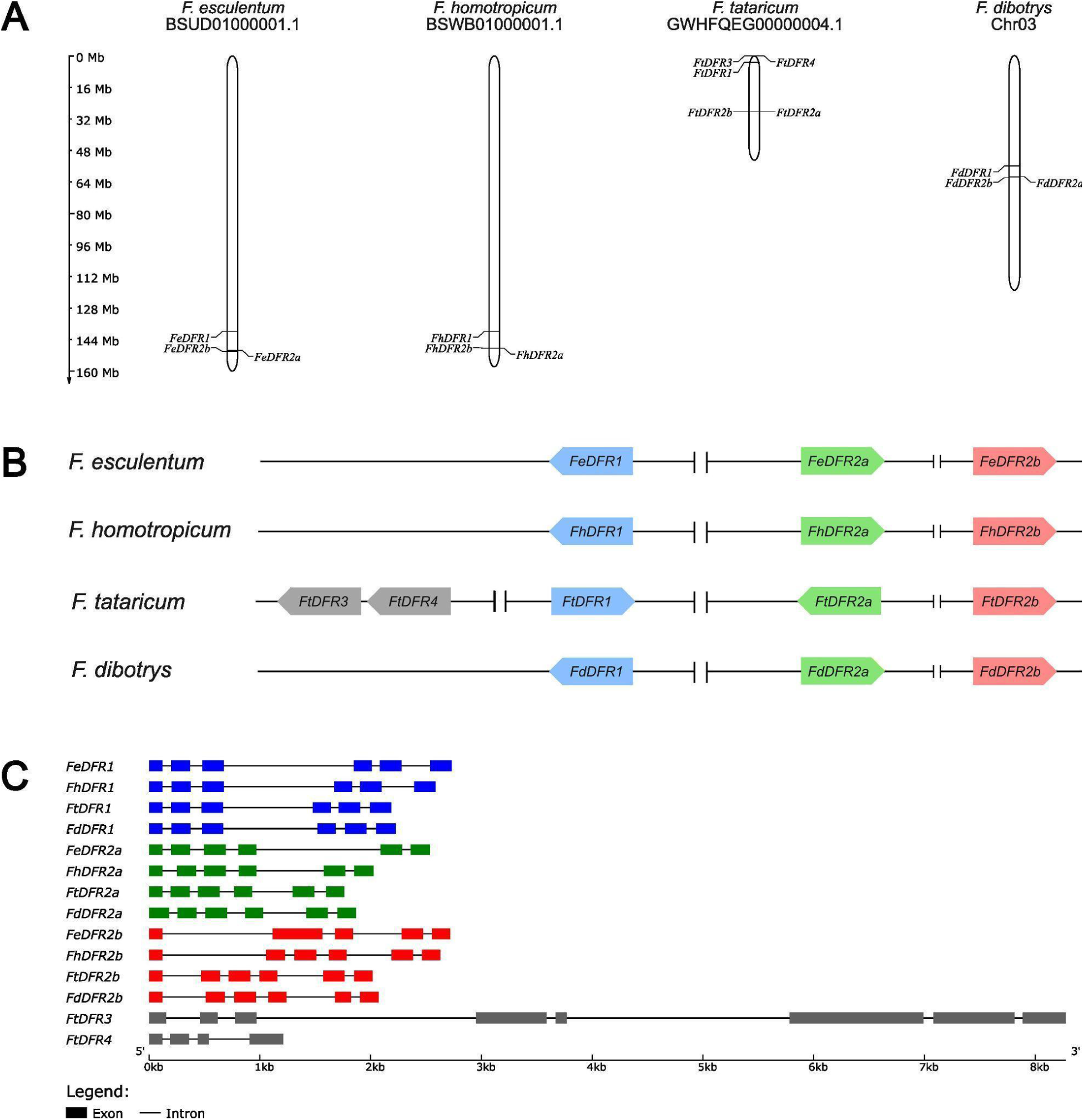
(A) Positions of each *DFR* duplicate in the chromosome and (B) their positions with arrow shape direction showing their orientation. (C) The exon and intron structure for each *DFR*. Blocks indicate the positions of exons, while lines indicate connecting introns. Different colors were used to highlight DFR1, DFR2a, DFR2b, and DFR3, respectively.

The *DFR* genes identified here were renamed based on their syntenic positions and placement in the constructed phylogenetic tree. They are referred to as *DFR1*, *DFR2a*, *DFR2b*, *DFR3*, and *DFR4*. *DFR2a* and *DFR2b* are located in close proximity, with a distance ranging from 40 Kbp in *F. dibotrys* to 400 Kbp in *F. esculentum* (**Figure 2A**). In contrast, *DFR1* is positioned at a distant locus of 4 to 27 Mbp away from the *DFR2* locus. This scattered pattern of gene duplication across the chromosome in *Fagopyrum* suggests a unique form of duplication that differs from the previously studied *DFR* duplications such as those reported in *Lotus japonicus*, where the *DFR* copies are arranged in tandem repeats [81]. The orientation of each duplicate is also conserved between orthologs across all *Fagopyrum* species with an exception of *F. tataricum* which is assumed to be caused by a secondary event such as an inversion (**Figure 2B**). *DFR1* in *Fagopyrum* maintains a conserved intron-exon structure, consisting of six exons and five introns, a pattern also observed in morning glories and onions [82, 83]. The *DFR2* genes generally have the same number of exons, except for *FeDFR2b* with five exons. In comparison, *FtDFR3* has eight exons and *FtDFR4* has four exons (Figure 2C).

#### Enzyme Similarities and Substrate Binding Region

Sequence similarities between the DFR proteins revealed a high degree of conservation within each ortholog group (**Figure 3A**). DFR1 share at least 99.4% similarity between species. Similarly, DFR2a and DFR2b show high similarity to their ortholog group with more than 91.9% for DFR2a and more than 90.7% for DFR2b. Notably, DFR2a and DFR2b are more similar to each other with at least 88.0% similar, than to *DFR1*, suggesting a closer evolutionary relationship. Interestingly, DFR3 and DFR4 in *F. tataricum* appear to be distinct from both DFR2 and DFR1 which strengthens the assumption that they might be pseudogenes.

**Figure 3:**
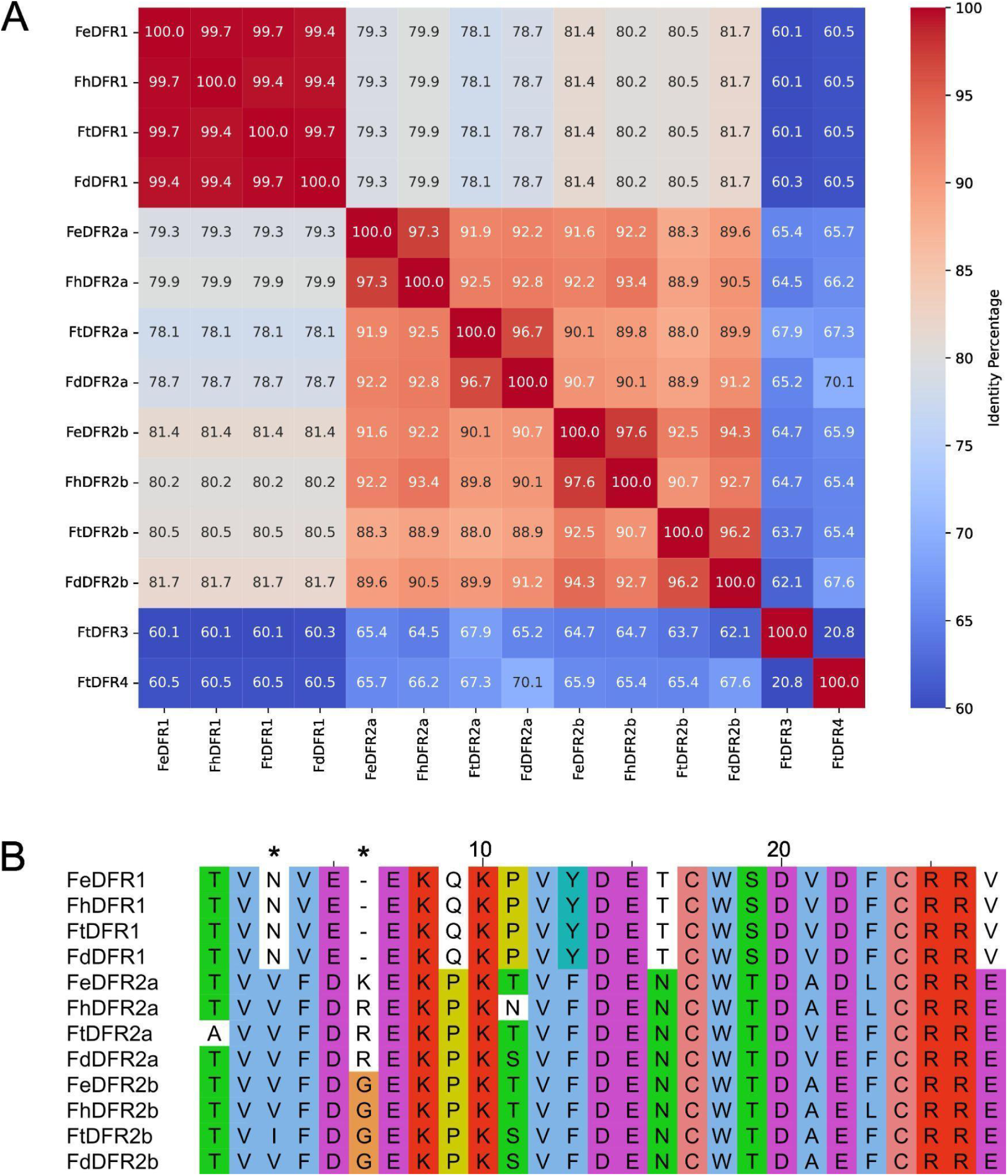
(A) Protein sequence similarities between each DFR. (B) The aligned sequences correspond to the 26-amino acid regions that determine substrate specificity. The first * on top of the alignment corresponds to the 3^rd^ position that is studied to be responsible for substrate specificity and the second * corresponds to the insertion in DFR2 duplicates between position 5 and 6. The full sequence alignment of each protein can be found in Additional file 2.

When the sequences were aligned, variations in the substrate binding domain were observed between DFR1 and DFR2 (**Figure 3B**). The substrate specificity is determined by a conserved region comprising 26 amino acids that has been well studied in other plants such as *Petunia*, *Gerbera* and *Cymbidium* [36, 37, 84, 85]. All DFR1 proteins contain an asparagine (N) at position 3, which permits acceptance of all three dihydroflavonols as substrates. In contrast, DFR2a and DFR2b show a substitution at this position, where all members have valine (V), except for FtDFR2b, which has isoleucine (I). This differs from other well-studied aspartic acid (D) substitution, which increases the preference for DHQ and DHM, and the alanine (A) substitution, which favours DHK while opposing DHM [34]. In addition, DFR2a and DFR2b contain a unique insertion at position 5-6, which is glycine (G) in all DFR2b, arginine (R) in FhDFR2a, FtDFR2a, FdDFR2a, and lysine (K) in FeDFR2a. This insertion is absent in DFR1 and other well-studied conserved DFR regions in plants. There exist also other substitutions in the conserved region of DFR2 genes when compared to DFR1, but they are still not yet known or studied for their consequences.

### Phylogeny of *DFR*

A tree classified *DFR* candidates from the buckwheat family by determining whether they were placed within the clade of known *DFR* genes in the buckwheat family or in the *DFR*-like clade. Some species within the family harbor at most one *DFR*-like gene. The classification of true *DFR* genes is further supported by comparison with other families in the *Caryophyllales*. Other species in the *Polygonaceae* family also exhibit multiple *DFR* duplications, particularly in the genera *Persicaria*. *Persicaria maculosa* has three *DFR* copies, while *Persicaria tinctoria* possesses four. *Fallopia convolvulus* contains two copies, whereas only one was identified in *Fallopia multiflora* and *Reynoutria japonica* has 3 *DFR* copies. All these *DFR* duplicates are found on different chromosomes, except for *R. japonica* that has two copies that are located on the same chromosome. In contrast, other species such as *Oxyria digyna*, the genus *Rheum* and the genus *Rumex* each have a single *DFR* copy. The *DFR* genes in *Fagopyrum* form two distinct clades. One clade harbours known *DFR* sequences from databases and is positioned near related species such as *Rheum*, *Oxyria*, *Polygonum*, and *Fallopia*. This group is designated as *DFR1*, as it is assumed to be the original copy that maintained the ancestral function (canonical *DFR*). In contrast, *Persicaria* species form a separate clade outside of the *Fagopyrum*-*Rheum*-*Polygonum* grouping, which is expected given their evolutionary divergence [86, 87]. *F. esculentum* is clustered closely with *F. homotropicum*, while *F. tataricum* is clustered closely with *F. dibotrys* (also known as *F. cymosum*), as they are closely related to each other, respectively. These four *Fagopyrum* species in this study are placed within the major cymosum group of the tribe *Fagopyreae*, in the subfamily *Polygonoideae* of the buckwheat family [39, 41]. An interesting finding is that the *DFR* duplicates in the *Fagopyrum* genus, including the *FeDFR2* sequences identified in a previous study [48], are positioned at the base of the *Polygonaceae* rather than clustering closely with *DFR* of other family members. This distinct clade, referred to as the *DFR2* clade, can be further divided into two subclades: *DFR2a* and *DFR2b*. Each *Fagopyrum* species possesses one *DFR* copy in each subclade except for *F. tataricum* with 3 *DFR* copies in *DFR2a* subclade (*FtDFR2a*, *FtDFR3*, and *FtDFR4*). *FeDFR1a* (LC216398.1) from Katsu et al. is clustered within the *DFR1* clade, while *FeDFR2* (LC216399.1) is clustered within the *DFR2a* clade (**Figure 4**).

**Figure 4:**
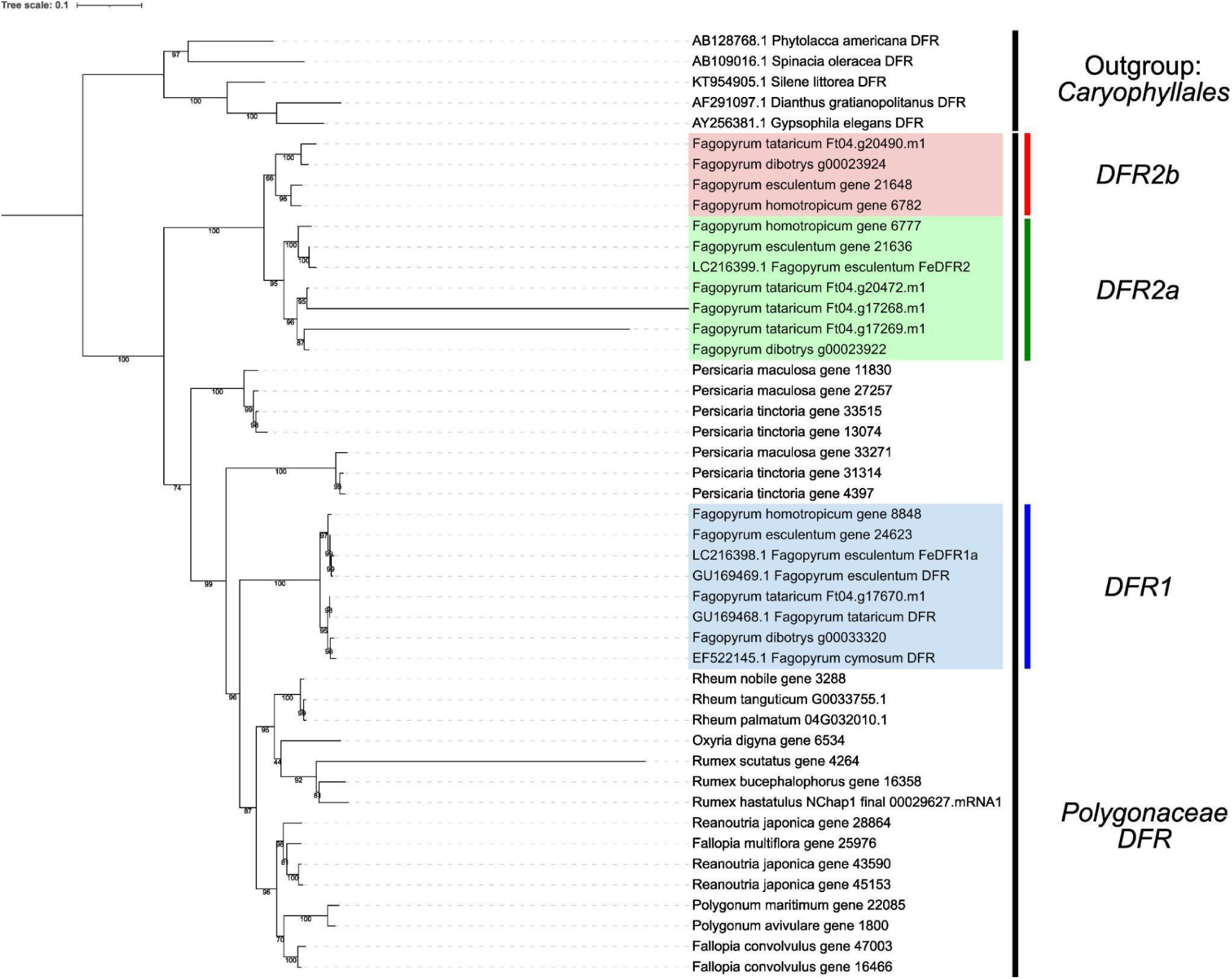
Maximum likelihood tree of *DFR* by IQ-TREE v2.4.0. Sequences retrieved from NCBI databases are labeled with their accession numbers and the other sequences were retrieved using KIPEs from each genome dataset. Sequences from the other family in the core *Caryophyllales* were used as outgroups. *DFR* from *Fagopyrum* are clustered into 3 main clades: *DFR1* (blue), *DFR2a* (green), and *DFR2b* (Red). See Additional file 3 for additional support.

### Synteny of *DFR*

Synteny analysis was performed using MCScan from JCVI to study the positions and order of genes on chromosomes across species, as well as to investigate their evolutionary origin. All four *Fagopyrum* genome sequences, together with those of closely related species such as *Rheum palmatum*, *Polygonum aviculare*, and *Fallopia multiflora*, were used to visualize the synteny of *DFR*. Macrosynteny analysis (**Figure 5A**) showed that the syntenic block harbouring *DFR1* is highly conserved among species of the family, with *DFR1* located in the middle of the block. This observation is also supported by the microsynteny analysis (**Figure 5B**). In contrast, the synteny of *DFR2* is more complex. There is no clear syntenic block that connects the *DFR2* locus in *Fagopyrum* with other family members. In macrosynteny, the *DFR2* locus is found outside of the syntenic blocks that harbour *DFR1* and lacks collinear or conserved flanking genes between *Fagopyrum* and other species. Interestingly, in *F. tataricum*, *DFR2a* and *DFR2b* do not follow this pattern. They are located at a different locus, which is instead replaced by *DFR3* and *DFR4*. Nevertheless, the names of the duplicates in *F. tataricum* remain as *FtDFR2a* and *FtDFR2b*, because their sequences are more similar (**Figure 3A**) and also closer to other *DFR2* orthologs in the phylogenetic tree (**Figure 4**). This difference in synteny pattern is likely due to another mechanism, most probably a chromosomal inversion that occurred after duplication during evolution. Microsynteny of *DFR2* (**Figure 5C**) further supports this, as the upstream region of *DFR2* genes is conserved across all species, but the region containing *DFR2* itself exists only in *Fagopyrum* and not in other family members.

**Figure 5:**
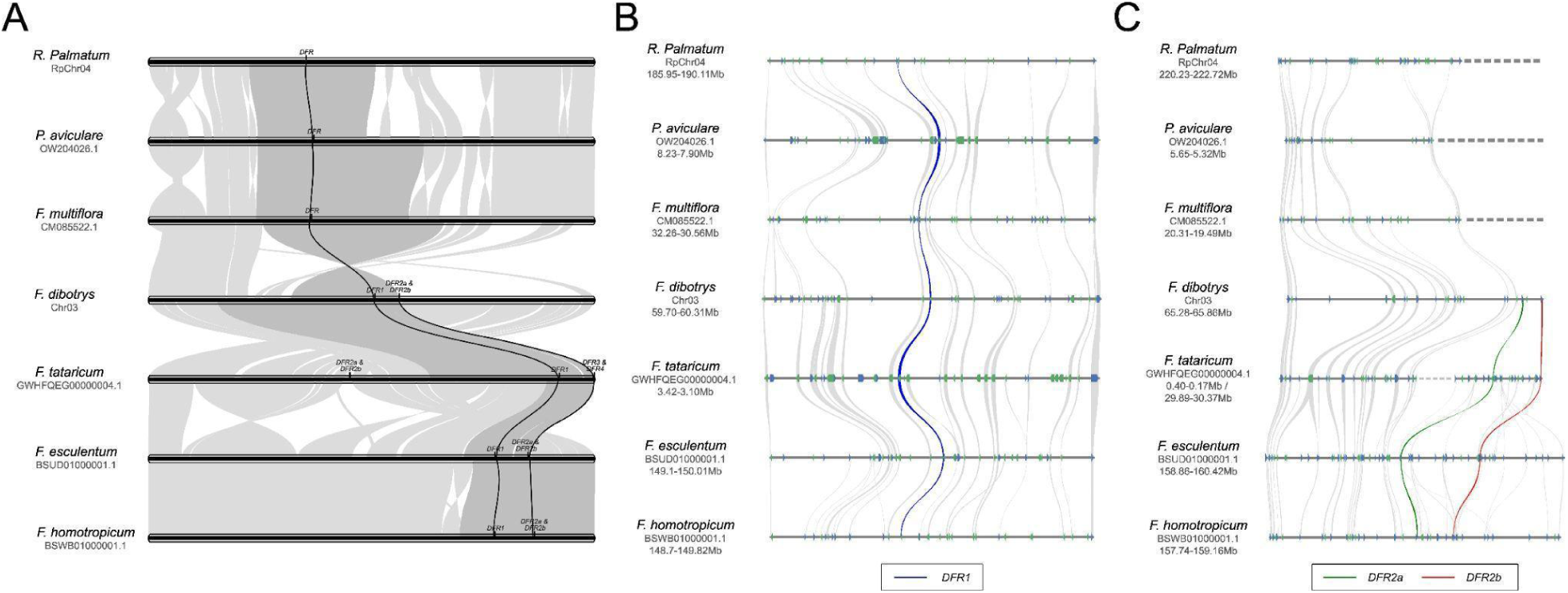
(A) Macrosynteny of the chromosome harboring *DFR* among *Fagopyrum* species and other species of buckwheat family *Rheum palmatum*, *Polygonum aviculare*, and *Fallopia multiflora*. The accession number of each chromosome is listed along with the species name. Black lines connecting species indicate the syntenic locus containing *DFR*, while gray waves represent other syntenic blocks between species. (B & C) Microsynteny of *DFR1* (blue), *DFR2a* (green), and *DFR2b* (red) among the species with the location of each locus in the chromosome displayed. The microsynteny of *DFR2* in *F. tataricum* is divided into two loci of different locations in its chromosome, divided by dotted lines. Dotted lines in *R. palmatum*, *P. aviculare*, and *F.* multiflora indicate that the genes *DFR2* and its flanking genes do not exist in these species.

### *DFR* Expression

RNA-seq datasets were retrieved from the SRA based on the availability of clear labels indicating the plant tissue of origin. For *F. esculentum* and *F. tataricum*, samples were available from root, leaf, flower, seed, stem, and seedling. For *F. dibotrys*, datasets were available for root, leaf, stem, rhizome, and tuber, but not for flower or seedling. *F. homotropicum*, on the other hand, had no RNA-seq datasets in the database. The SRA run IDs for each species, grouped by tissue, are listed in Additional file 4. Boxplots were generated to visualize transcript abundance in transcripts per million (TPM) for each gene (**Figure 6**). Across all species, *DFR1* consistently shows the highest expression across tissues. In *F. esculentum*, *DFR1* expression is strongest in the root, with a median TPM of ∼600, while other tissues range between 200–300 TPM. In *F. tataricum*, the highest expression is found in the flower (∼200 TPM), followed by the root (∼170 TPM). In *F. dibotrys*, which had fewer RNA-seq datasets, the highest expression is observed in the tuber (∼600 TPM), with other tissues showing lower expression between 50–200 TPM. In contrast, *DFR2a* and *DFR2b* show divergent and generally lower expression compared to *DFR1*. In *F. esculentum*, only *FeDFR2a* is expressed, mainly in root and seed, with values below 50 TPM. In *F. tataricum*, both *FtDFR2a* and *FtDFR2b* are expressed, with *FtDFR2a* showing relatively high expression in seed, and *FtDFR2b* in root. *FtDFR4* shows low expression in some tissues, but this is likely due to background noise. In *F. dibotrys*, both *FdDFR2a* and *FdDFR2b* show no noticeable expression in any tissue.

**Figure 6:**
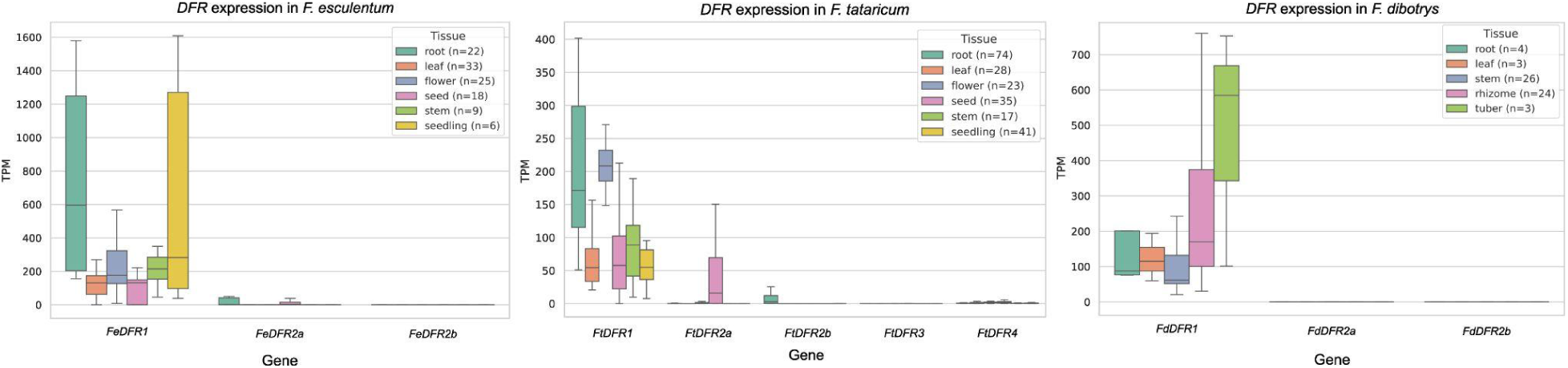
Boxplot diagrams illustrating the expression levels of *DFR* duplicates in *F. esculentum*, *F. tataricum*, and *F. dibotrys*. TPM = transcripts per million.

### Cis-regulatory Element

To further investigate the expression differences between *DFR* copies, promoter sequences were analyzed for major transcription factor binding motifs. The search focused on MYB and bHLH binding sites, specifically: MYBCORE (CNGTTR), MYBPLANT (MACCWAMC), MYBPZM (CCWACC), and MYCCONSENSUS (also known as the E-box or bHLH binding site, CANNTG). MYBCORE, MYBPLANT, and MYBPZM are proposed MYB transcription factor binding sites [88–91], while MYCCONSENSUS is recognized as a bHLH binding site [92]. These motifs are crucial for MYB and bHLH transcription factors to bind and form the MBW complex with WD40, which regulates *DFR* expression [38, 93]. The analysis revealed an abundance and diverse pattern of MYB and bHLH recognition elements in the promoter regions of *DFR* genes across all species (**Figure 7**). In particular, several MYB binding sites are conserved across all *DFR1* copies, including two MYBPLANT and one MYBPZM element located within the proximal 400 bp upstream of the putative TSS. MYBCORE sites are more dispersed throughout the promoter but show partial conservation in the *FeDFR1* and *FhDFR1* promoters, likely reflecting their close relationship. The MYCCONSENSUS (bHLH) binding site located ∼200 bp upstream of the TSS is conserved across all *DFR1* promoter sequences. Three such bHLH sites were identified, two of which are located close together, possibly facilitating bHLH transcription factor dimerization [94]. Beyond the 400 bp region upstream of the TSS, binding sites appear more randomly and are not conserved across species. In contrast, *DFR2* promoters generally lack MYB binding sites, especially within the proximal 400 bp. An exception is *FtDFR2a*, which contains one MYBCORE and two MYBPLANT motifs. *FeDFR2b* and *FdDFR2a* each contain two MYBCORE motifs located around -400 bp. Despite the overall scarcity of MYB sites, a conserved pair of bHLH binding sites is present in nearly all *DFR2a* and *DFR2b* promoters, except for *FtDFR2a*, *FtDFR3*, and *FtDFR4*. These conserved bHLH sites may have originated from those found in *DFR1* and could still support gene expression, even in the absence of MYB motifs.

**Figure 7:**
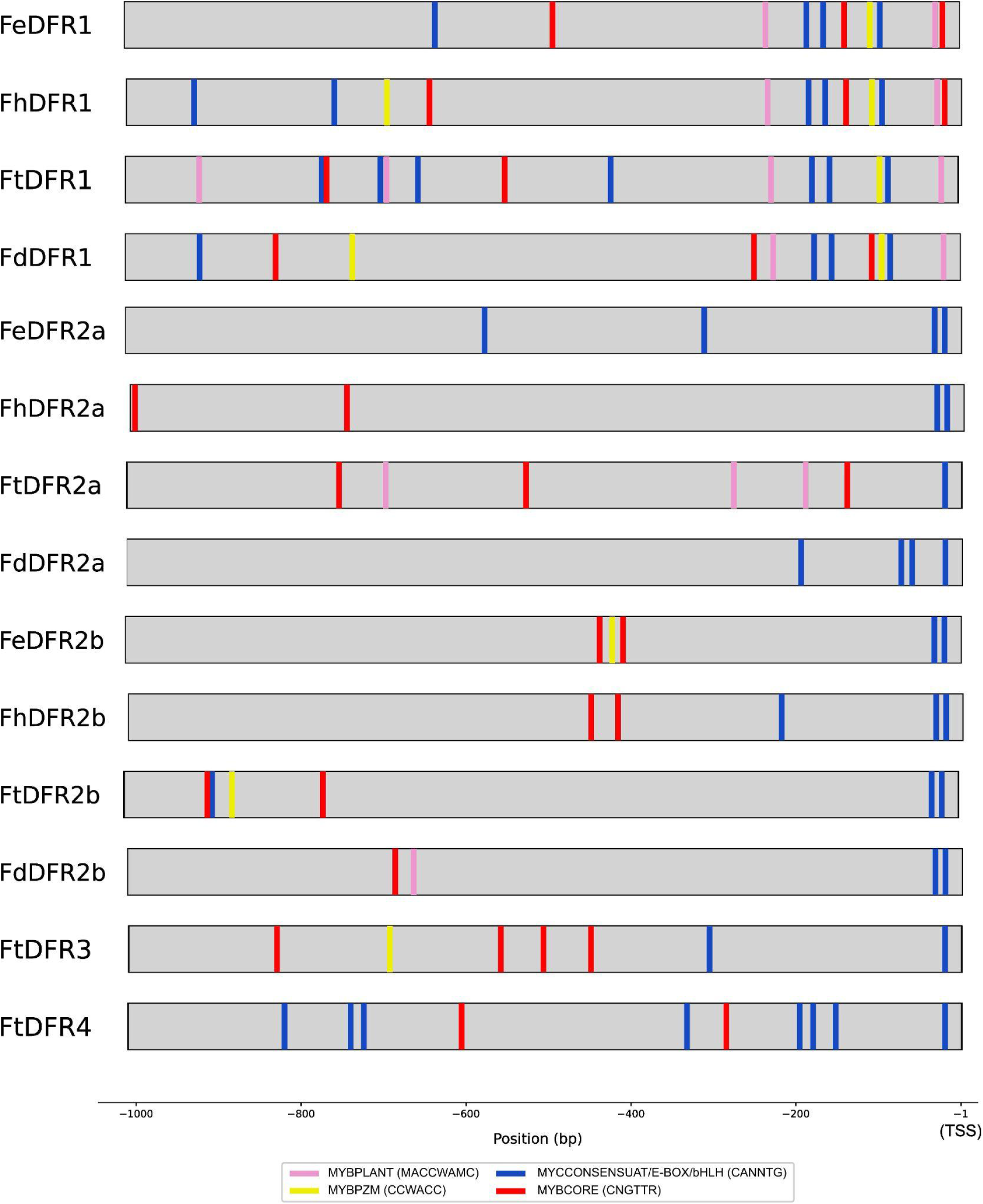
Schematic representation of transcription factor binding motifs in the promoter regions. The putative transcription start site (TSS), identified based on RNA-seq alignment to the genome, is set as position +1.

### Selective Pressure Analysis

To examine the selection pressures acting on the *DFR* duplicates, the Ka and Ks values were calculated for each orthologous and paralogous gene pair within the genus *Fagopyrum*. Gene pairings were determined based on their orthologous partners between species (Table 1) and their paralogous pairs within the same species (Table 2). This analysis was performed on both the entire *DFR* coding sequence and on the conserved 26–amino acid region to assess whether selection pressures differ between the two regions (Table 1, Table 2, Additional file 5).

**Table 1:** Comparison of substitution rates between orthologous *DFR* duplicates among *Fagopyrum* species. The comparison is done for the whole coding sequence (CDS) and their corresponding 26-amino acid conserved region. The full table can be found in Additional file 5.

| Sequence | Full CDS |  | Conserved Region |  |
| --- | --- | --- | --- | --- |
|  | Ka/Ks | P-Value (Fisher) | Ka/Ks | P-Value (Fisher) |
| *FeDFR1_FhDFR1* | 0.042 | 0.0 | NA | NA |
| *FeDFR1_FtDFR1* | 0.009 | 0.0 | 0.001 | 0.0 |
| *FeDFR1_FdDFR1* | 0.011 | 0.0 | 0.001 | 0.003 |
| *FhDFR1_FtDFR1* | 0.022 | 0.0 | 0.001 | 0.0 |
| *FhDFR1_FdDFR1* | 0.011 | 0.0 | 0.001 | 0.003 |
| *FtDFR1_FdDFR1* | 0.001 | 0.0 | 0.001 | 0.080 |
| *FeDFR2a_FhDFR2a* | 0.070 | 0.0 | 50.0 | 0.321 |
| *FeDFR2a_FtDFR2a* | 0.174 | 0.0 | 0.220 | 0.016 |
| *FeDFR2a_FdDFR2a* | 0.179 | 0.0 | 0.166 | 0.0 |
| *FhDFR2a_FtDFR2a* | 0.142 | 0.0 | 0.261 | 0.032 |
| *FhDFR2a_FdDFR2a* | 0.146 | 0.0 | 0.188 | 0.008 |
| *FtDFR2a_FdDFR2a* | 0.184 | 0.0 | 0.184 | 0.047 |
| *FeDFR2b_FhDFR2b* | 0.143 | 0.0 | 1.029 | 0.683 |
| *FeDFR2b_FtDFR2b* | 0.098 | 0.0 | 0.056 | 0.0 |
| *FeDFR2b_FdDFR2b* | 0.094 | 0.0 | 0.002 | 0.0 |
| *FhDFR2b_FtDFR2b* | 0.135 | 0.0 | 0.109 | 0.001 |
| *FhDFR2b_FdDFR2b* | 0.157 | 0.0 | 0.059 | 0.0 |
| *FtDFR2b_FdDFR2b* | 0.202 | 0.0 | 0.528 | 0.199 |

**Table 2:** Comparison of substitution rate between paralogous *DFR* duplicates among *Fagopyrum* species. The comparison is done for the whole coding sequence (CDS) and their corresponding 26-amino acid conserved region. The full table can be found in Additional file 5.

| Sequence | Full CDS |  | Conserved Region |  |
| --- | --- | --- | --- | --- |
|  | Ka/Ks | P-Value (Fisher) | Ka/Ks | P-Value (Fisher) |
| *FeDFR1_FeDFR2a* | 0.022 | 0.0 | 0.059 | 0.0 |
| *FeDFR1_FeDFR2b* | 0.030 | 0.0 | 0.059 | 0.0 |
| *FeDFR2a_FeDFR2b* | 0.136 | 0.0 | 1.787 | 0.906 |
| *FhDFR1_FhDFR2a* | 0.021 | 0.0 | 0.076 | 0.0 |
| *FhDFR1_FhDFR2b* | 0.040 | 0.0 | 0.066 | 0.0 |
| *FhDFR2a_FhDFR2b* | 0.113 | 0.0 | 50.0 | 0.682 |
| *FtDFR1_FtDFR2a* | 0.045 | 0.0 | 0.065 | 0.0 |
| *FtDFR1_FtDFR2b* | 0.031 | 0.0 | 0.053 | 0.0 |
| *FtDFR2a_FtDFR2b* | 0.181 | 0.0 | 0.616 | 0.394 |
| *FdDFR1_FdDFR2a* | 0.040 | 0.0 | 0.056 | 0.0 |
| *FdDFR1_FdDFR2b* | 0.030 | 0.0 | 0.066 | 0.0 |
| *FdDFR2b_FdDFR2b* | 0.167 | 0.0 | 0.175 | 0.045 |

#### Orthologous Pair

Table 1 shows that *DFR1* orthologs are generally well conserved, both across their entire sequence and in the conserved region. The Ka/Ks values were below 0.04 for the full coding sequence and below 0.003 for the conserved region. *F. esculentum* and *F. homotropicum* show few substitutions (Additional file 5) when compared, as do *F. tataricum* and *F. dibotrys*, which are expected given their close phylogenetic relationships. Notably, the conserved region of *DFR1* in *F. esculentum* and *F. homotropicum* shows no substitutions at all. The number of nonsynonymous substitutions is also low, with a maximum of two for the full gene and none in the conserved region. In contrast, *DFR2a* orthologs show generally higher Ka/Ks values than *DFR1*. For example, comparison between *FeDFR2a* and *FhDFR2a* gives a Ka/Ks value of 0.07 for the full CDS but an unusually high value of 50 for the conserved region, because all four substitutions in this region are nonsynonymous. However, this small number of substitutions also results in a higher p-value, making the result less reliable. Other pairwise comparisons of *DFR2a* and *DFR2b* orthologs generally show similar Ka/Ks values for both the full CDS and the conserved region, with the exception of *FeDFR2b* vs. *FhDFR2b* and *FtDFR2b* vs. *FdDFR2b*, where the conserved region shows higher values.

#### Paralogous Pair

Across all paralogous comparisons in Table 2, *DFR1* consistently shows low Ka/Ks ratios when compared to *DFR2a* or *DFR2b*. Specifically, Ka/Ks values remain below 0.05 for the full CDS and below 0.08 for the conserved region. The total number of substitutions is also relatively consistent, ranging from 218–242 in the full CDS, and 28–31 in the conserved region. Another clear trend is that the number of substitutions between *DFR1* and *DFR2b* is lower than between *DFR1* and *DFR2a* in all species. In contrast, comparisons between *DFR2a* and *DFR2b* show higher Ka/Ks values, though still below 0.5 for the full CDS. In the conserved region, however, the ratios can be much higher, for example, 1.787 for *FeDFR2a* vs. *FeDFR2b* and 50 for *FhDFR2a* vs. *FhDFR2b*. Interestingly, comparisons of *FtDFR2a* vs. *FtDFR2b* and *FdDFR2a* vs. *FdDFR2b* show lower Ka/Ks values for the conserved region, only 0.616 and 0.175, respectively.

## Discussion

### Hypothetical Mechanism of Duplication

It can be assumed that *DFR1* represents the gene copy with the original ancestral function, while *DFR2* genes are more recent duplicates that are diverging from the main function of bona fide *DFR*. This assumption is supported by three lines of evidence. First, *DFR1* is located in a conserved syntenic region with *DFR* genes in other plant species, both within the *Polygonaceae* family and the broader *Caryophyllales* order. Second, *DFR1* consistently shows the highest expression levels compared to the other duplicates, strongly suggesting that it retains the ancestral function, whereas the duplicates with low or no expression may be pseudogenes or in the process of pseudogenization. Third, *DFR1* clusters phylogenetically closer with *DFR* genes from related species (**Figure 4**). The observation that *DFR2a* and *DFR2b* are located at different genomic loci than *DFR1* raises questions about the mechanism of their origin, since this pattern does not fit the typical case of gene duplication. WGD was considered as a possible origin for *DFR2*, however, comparisons with other *Polygonaceae* species indicate that *DFR* duplicates often reside on different chromosomes. Chromosome number variation and polyploidy patterns in *Fallopia* and *Persicaria* species suggest historical WGDs in some taxa [95–97], but the genomic distribution of *DFR2* in *Fagopyrum* does not support a WGD origin. Transposable element (TE)-mediated duplication was also considered, given the high TE content and evidence of recent TE proliferation in *F. esculentum* [11, 98]. While TEs are present near *DFR1* and *DFR2* loci, the retention of introns in *DFR2a* and *DFR2b* rules out retrotransposon-mediated duplication. TE annotations using EDTA [78] show no TIR or DNA transposons including helitrons that directly flank the *DFR* duplicates, suggesting that TEs may not have directly mediated the duplication [13]. Segmental duplication was further evaluated using dot plot analyses through ModDotPlot [79] (Additional file 6). No patterns indicative of segmental duplication was observed between *DFR1* and *DFR2* loci in *Fagopyrum*, although repetitive sequences are present near both loci. These repeats may represent TEs or other repetitive elements, but they do not appear sufficient to explain the duplication mechanism. Therefore, while *DFR2* likely arose through a mechanism independent of WGD or retrotransposition, the exact process potentially involving TE-associated rearrangements remains unclear.

#### DNA Duplication Associated with Inversion

This case of duplication from *DFR1* to *DFR2* appears to deviate from typical duplication mechanisms and was therefore examined for alternative explanations. Several clues support this: first, the *DFR2* copies are assumed to be located at the breakpoints of genome inversions or rearrangements (**Figure 5A**); second, the orientation of *DFR2* is inverted relative to *DFR1* and the two loci are separated by a considerable genomic distance (**Figure 2**). Additional observations include frequent genome rearrangements both within *Fagopyrum* species and between *Fagopyreae* and other members of the *Polygonaceae* family. These patterns suggest that the *DFR* gene duplications may be associated with structural variations, such as inversions, rather than arising from tandem or segmental duplications. These observations point towards a mechanism of gene duplication associated with chromosomal inversions, first described by Ranz et al. [99] in the genome of *Drosophila melanogaster*, and later observed in other species such as *Drosophila buzzatii* [100], *Drosophila mojavensis* [101] and even in the bacterium *Helicobacter pylori* [102]. This mechanism has also been discussed in a review on chromosomal inversions by Casals and Navarro [103] that differentiate between ectopic recombination and this staggered break model. Furuta et al. referred to this process as “DNA duplication associated with inversion” (DDAI), a term that will be used from this point on in this study [102]. According to them, DDAI involves the duplication of a DNA segment at one chromosomal locus and its reinsertion at a distant site on the same chromosome in an inverted orientation, accompanied by an inversion of the sequence in between [102]. Ranz et al. described it as an inverted duplication associated with inversion breakpoint regions resulting from staggered breaks [99]. The mechanism of this type of gene duplication will serve as the working hypothesis for the duplication of *DFR1* to *DFR2* in this study (**Figure 8**). In this model, a segment of DNA on a chromosome undergoes two pairs of staggered single-strand breaks at its ends, resulting in long 5’ overhangs. The segment is then inverted and joined into the opposite breakpoints. The breaks are then repaired by non-homologous end joining (NHEJ), in which the 5’ overhangs are filled in and ligated to restore DNA continuity through replication. If a gene lies within the region affected by the staggered break and is successfully retained during inversion and repair, this results in a duplicated copy of the gene in an inverted orientation relative to the ancestral copy - without disrupting the original gene. A possible alternative to this mechanism involves only one pair of staggered single-strand breaks at one end and a double-strand break at the other. A hallmark of this mechanism is that gene copies are found in an inverted orientation, near chromosomal inversion breakpoints, and are present only in species that have the specific chromosomal rearrangement. Using both experimental and computational analysis, Ranz et al. found that 17 out of 29 chromosomal inversion breakpoint regions between *D. melanogaster* and its close relatives *D. simulans* and *D. yakuba* are associated with inverted gene duplications resulting from these inversions [99]. The average nucleotide identity (± standard deviation) between the duplicated gene pairs was approximately 88% ± 5.4%, which closely resembles the similarity observed in this study (**Figure 3A**). In the case of the *DFR2* duplication, all the key features supporting the DDAI mechanism are present. The formation of *DFR2a* and *DFR2b* can be further explained as resulting from a subsequent tandem or proximal duplication of the initial *DFR2* copy generated through DDAI (**Figure 8E,F**). A comparable example is observed in *D. melanogaster*, where the genes *CG31286* and *CG34034*, located at two separate breakpoints, underwent tandem duplication following a DDAI event associated with inversion 3R(7) when compared to *D. simulans* [99]. Furuta et al. studied the “birth” and “death” of genes linked to chromosomal inversions in *H. pylori*, a bacterium with high genome plasticity. By comparing multiple genomes, they found that gene gains and losses often occurred at inversion breakpoints, providing evidence for DNA duplication associated with inversion (DDAI). They reported cases of complete and inverted gene duplication, partial duplication, and gene decay. Importantly, they showed that the near-identical homology created by DDAI events promotes subsequent homologous recombination, although this recombination decreases as sequence divergence accumulates over time [102]. Calvete et al. analyzed the 2m and 2n inversions in *D. buzzatii*. Inversion 2m was associated with a 13 kb duplicated segment generated by staggered single-strand breaks and repair via NHEJ. The segment contained five genes, but some copies became pseudogenes or were lost. Inversion 2n was instead linked to double-strand breaks and NHEJ repair, and both inversions may have shared breakpoints [100]. Guillén and Ruiz reported two inversions in *D. mojavensis* (2h and 2q), both involving inverted duplications of non-repetitive DNA created by staggered single-strand breaks and NHEJ repair. The 2h inversion contained a 7.1 kb segment with three genes, while 2q involved a smaller 1 kb duplicated segment [101].

**Figure 8:**
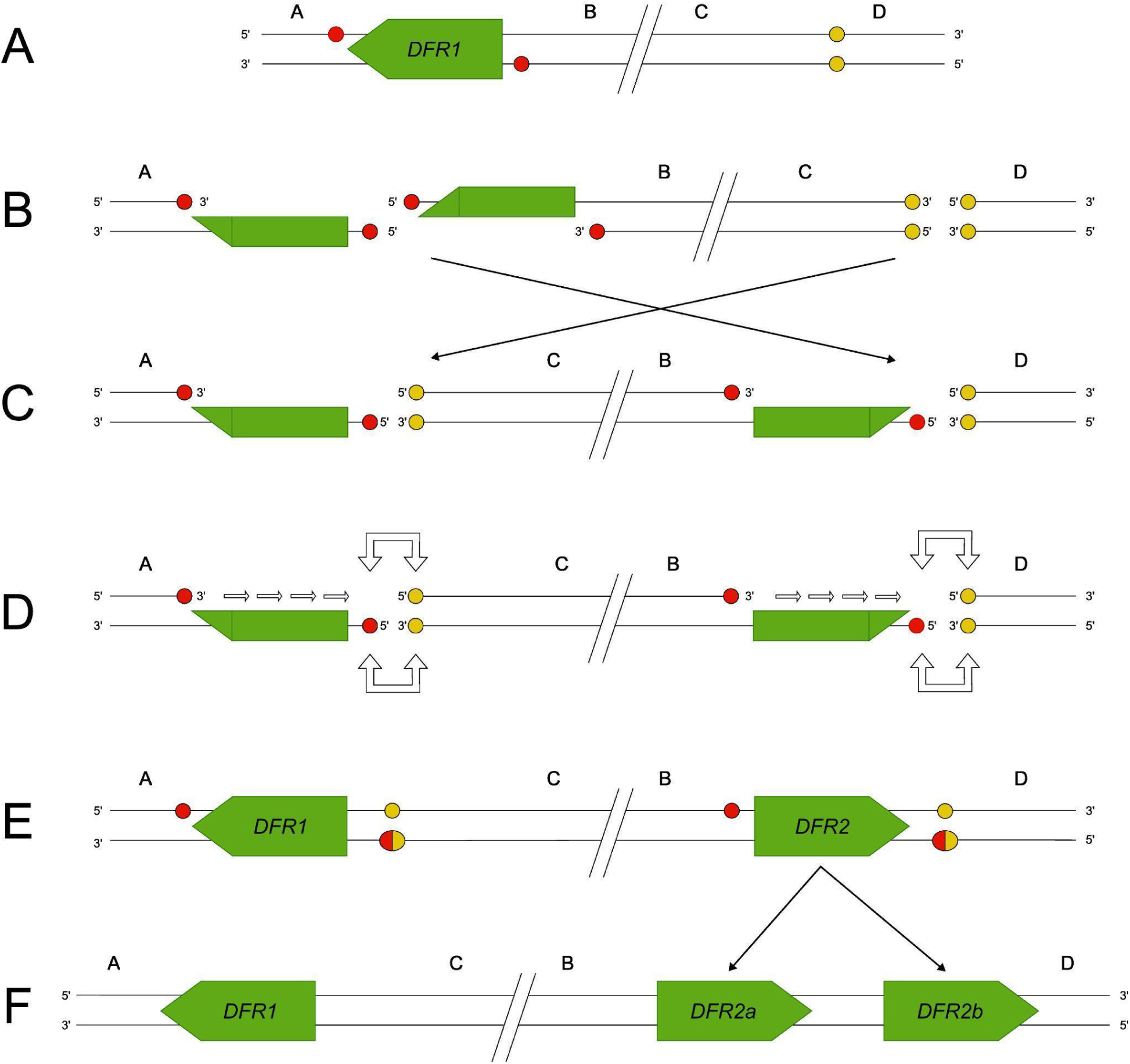
Proposed mechanism of *DFR* duplication in *Fagopyrum*. (A) The hypothetical ancestral state in the common ancestor of *Fagopyrum* and its family members. (B) A pair of staggered single-strand breaks, containing the *DFR* sequence, results in a long 5’-overhang at one end of the segment, with a double-strand break at the other end. (C) The segment is inverted, and (D) the 5’-overhang is filled and the end of the segments are ligated to the corresponding DNA segments by the NHEJ repair mechanism (depicted with white arrows). (E) The intermediate state following the repair process, and (F) the tandem or proximal gene duplication event leading to *DFR2* being duplicated into *DFR2a* and *DFR2b*.

#### Preexisting Gene Duplications at Breakpoints

An important finding in the study of Guillén and Ruiz concerns the presence of preexisting gene duplications at inversion breakpoints [101]. They observed four cases where inversion breakpoints were located between preexisting duplicated genes. Their statistical analysis showed that the breakpoints were significantly more likely to be located between duplicated genes than would be expected by chance. Guillén and Ruiz proposed two possible explanations for this observation. First, they suggested that duplicated genes might cause genomic instability, leading to an increased rate of double-strand breaks (DSBs) or may create “permissive” regions where breakage is more likely. Alternatively, they hypothesized that the mobilization of a duplicated gene could provide beneficial positional effects, supporting in the fixation of the inversion within the species. When duplicated genes are co-located in the same chromosomal domain, their evolution may be constrained due to the shared regulatory sequences, and sequence homogenization by frequent conversion or ectopic recombination events [101]. Relocating one of the copies to a different chromosomal region could relieve these constraints and potentially lead to advantageous regulatory changes [101], which could also be the case for *DFR1* and *DFR2*. In a comparative study between *Drosophila arizonae* and *D. mojavensis*, Runcie and Noor identified two inversion breakpoints that were located near an inverted gene duplication and a common repetitive element [104]. However, they found that the inverted gene duplication likely predated the inversion event and therefore rejected the DDAI-like hypotheses.

The duplication was hypothesized to have originally been present in the non-inverted ancestral chromosome, although the ancestral copy appeared to have degenerated [104]. In this study, it remains unclear whether the inverted gene duplication predated the inversion, as no dataset from a primitive *Fagopyrum* species or a most recent common ancestor without the inversion is available. This finding presents an alternative hypothesis for the *DFR2* duplication in *Fagopyrum*. In this scenario, a tandem gene duplication likely occurred first, leading from *DFR1* to *DFR2*. The inversion event would have occurred later, resulting in the original or its derived copies being translocated to a distant locus by chromosomal inversion resulting in an inverted orientation. The further tandem gene duplications of *DFR2* into *DFR2a* and *DFR2b* could have occurred either before or after the inversion event.

### Evolutionary Implications of *DFR* Gene Duplications

#### Duplication Mechanism and Timing

The duplication of *DFR* genes in *Fagopyrum* species is hypothesized to have occurred via DDAI, followed by a tandem gene duplication, resulting in three *DFR* copies. An alternative hypothesis would be that a tandem gene duplication occurred first, followed by an inversion that relocated one copy to a distant locus and another gene duplication resulting in *DFR2a* and *DFR2b*. The origin of *FtDFR3* and *FtDFR4*, however, remains unclear. It is assumed to have arisen via a DDAI event, as it is located at a putative chromosomal breakpoint (**Figure 5**). Notably, the intermediate chromosomal state between the ancestral synteny and the current arrangement following the hypothesized DDAI event has not yet been verified. Genome sequences of only four *Fagopyrum* species have been assembled to date, and all belong to the same *cymosum* subgroup within the *Fagopyreae* tribe and no genomes from the *urophyllum* subgroup have yet been sequenced. Extending the genome assembly of more species in both subgroups would be very informative to search for tracing the origin and evolutionary history of the *DFR* duplication. Nonetheless, evidence for chromosomal inversions involving the *DFR* locus in *Fagopyrum* exists. When macrosynteny between *Fagopyrum* and other *Polygonaceae* members (e.g., *F. multiflora* and *P. aviculare*) is examined, *DFR2a* and *DFR2b* in *F. dibotrys* are found near one end of a chromosomal breakpoint (**Figure 5**). This breakpoint may have been reused again during the DDAI event that duplicated *DFR*. It is also possible that multiple inversions occurred after the initial DDAI event, potentially driven by homologous recombination between regions of high sequence identity generated during the duplication process [102]. The exact breakpoints responsible for the duplication and inversion remain unidentified and require further investigation. However, dot plot comparisons between the duplicated loci (Additional file 6) reveal homologous regions that may result from the presence of TEs. These elements could contribute to the sequence similarity necessary to facilitate recombination, further supporting the possibilities of ectopic recombinations involving the syntelog containing *DFR* following the DDAI [102]. The gene sequence similarity, structural conservation, substrate-binding regions, and phylogenetic placement all confirm that the observed gene copies are indeed authentic DFRs, rather than unrelated genes or pseudogenes mistakenly identified. However, there remains ambiguity regarding the phylogenetic placement of the *DFR2* copies. In the constructed phylogenetic tree, they cluster near the base of the *Polygonaceae* family (**Figure 4**), rather than grouping closely with the known clade of *Fagopyrum DFR1*. This suggests that the duplication event leading to *DFR2* may have occurred before the divergence of species within the family. However, this scenario is less parsimonious. If the duplication had occurred prior to the diversification of the *Polygonaceae* family, one would expect other family members to retain at least one additional *DFR* copy resulting from the same duplication event. This is, however, not observed except for *R. japonica* that has 2 *DFR* duplicates on a single chromosome.

#### The Fate of *Fagopyrum DFR* Duplicates

The 26-amino-acid region responsible for substrate specificity [37] displays distinct features in *DFR2a* and *DFR2b*, with all duplicates containing valine at the third position, except for *FtDFR2b*, which contains isoleucine. Genes with valine at this position have been reported to have high affinity toward DHQ over other dihydroflavonols [48, 105] but the functional implications of isoleucine at this site in *FtDFR2b* remain unknown. Additionally, all derivative *DFR* genes possess an extra amino acid between positions 5 and 6 (**Figure 3B**), which is glycine in all *DFR2b*, arginine in *DFR2a*, except in *FeDFR2a* with lysine. The insertion of glycine in *FeDFR2* is hypothesised by Katsu et al. to increase the flexibility and alter the conformation of the loop spanning residues 4 to 18, which overlays the catechol ring of DHQ [48]. Although none of the residues in this loop directly interact with the substrate, such conformational changes may influence substrate selection [48]. Their docking models of the two proteins in complex with the three substrates suggested that *FeDFR1a*, with asparagine at the third position, could form more hydrogen bonds with each substrate than *FeDFR2*, which has valine at the same site [48]. The steric hindrance from a bulky Phe133 residue in *FeDFR2* may account for its preferential activity toward DHQ over DHM [48] and thus, both the identity of the amino acid at the third position and the surrounding residues forming the substrate-binding pocket contribute significantly to the substrate specificity of *DFR* enzymes in *F. esculentum*. The functional impact of the alternative inserted residues (arginine or lysine) at this position remains unclear and requires further investigation. The sequence divergences suggest that the duplicated *DFR* genes have undergone structural evolution in their substrate-binding regions, potentially altering their functional roles in flavonoid biosynthesis. Differential gene expression data indicate that some *DFR2* genes are only expressed in seeds and roots, whereas *DFR1* is broadly expressed across tissues. This is not unusual among duplicates in other organisms, as approximately 70% of gene pairs in *Arabidopsis* exhibit significant differences at the transcriptional level [2]. This tissue-specific expression pattern likely results from differences in cis-regulatory elements in their promoters. Notably, the promoter regions of *DFR2* contain fewer MYB-binding sites and predominantly feature bHLH-binding motifs, which may also explain the overall lower expression levels of *DFR2* [38]. This is also supported by the promoter analysis from Katsu et al. [48], that *FeDFR2* lacks the MYBPLANT and MYBPZM motifs, known binding sites for MYB transcription factors. Their gene expression analysis via real-time PCR across six organs (roots, stems, leaves, buds, flowers, and seeds) showed that *FeDFR1a* was expressed in all examined tissues while *FeDFR2* was predominantly expressed in roots and seeds, with expression levels in these organs approximately two orders of magnitude higher than those of *FeDFR1a* [48]. In contrast, our analysis shows *DFR2* expression is lower than *DFR1* (**Figure 6**), likely due to methodological differences. Further research is needed to clarify the regulatory elements driving tissue specificity, the DHQ preference of DFR2, and the functional effects of the different amino acid insertions.

Selective pressure analysis supports the functional relevance of the *DFR* copies. Orthologous comparisons of *DFR1* across species show very low Ka/Ks values, indicating strong purifying selection and conservation as the primary functional *DFR* gene in *Fagopyrum*. *DFR2a* and *DFR2b* orthologs exhibit slightly higher, yet still <0.5 Ka/Ks values, suggesting that they too are under purifying selection after the duplication event, albeit slightly relaxed. Paralogous comparisons reveal that *DFR2a* and *DFR2b* versus *DFR1* have Ka/Ks values below 0.1, indicating strong purifying selection and retention of *DFR* structure. Interestingly, when *DFR2a* and *DFR2b* are compared with each other, higher Ka/Ks values are observed, particularly within conserved regions, implying divergence in different functional directions. This could suggest that while both copies maintain the ancestral sequence and structure, they may be evolving distinct roles to reduce functional redundancy. The absence of pseudogenization or genetic drift features (Ka/Ks ≈ 1) in any of the pairwise comparisons further support the idea that all *DFR* copies remain under selective pressure and are likely beneficial to the genome. After gene duplication, the rate of sequence evolution is generally expected to increase, at least initially, due to relaxed selection against previously deleterious mutations. Consistent with this, the *DFR* duplicates exhibit relaxed purifying selection and sequence divergence. However, this increase in evolutionary rate is not always symmetric between duplicates [2], as shown here where *DFR2* exhibits a higher mutation rate than the ancestral *DFR1*. The most common fate of a duplicated gene is partial or complete deletion, reverting the locus to a singleton state [15]. Duplications that are retained may diverge to provide new functions (neofunctionalization) or split the ancestral function between copies (subfunctionalization). Whether the case of *DFR* duplicates represents neofunctionalization or subfunctionalization remains unclear, as changes in expression levels or tissue specificity alone do not conclusively determine either outcome [106]. Subfunctionalization or duplication-degeneration-complementation (DDC) involves the retention of both duplicates to preserve the full spectrum of ancestral functions which is here the ability to accept all three dihydroflavonols. A possible explanation is that after *DFR* duplication, each copy randomly lost part of its functional capacity (degeneration), and because the losses affected different subfunctions, both duplicates were retained to complement each other. Subfunctionalization does not always partition ancestral functions symmetrically, as one copy typically retains significantly more subfunctions than its paralog [2]. This aligns with the findings of Katsu et al., who showed that *FeDFR1a* has higher activity toward DHM and DHK but lower efficiency for DHQ, while *FeDFR2* shows high catalytic efficiency toward DHQ but lower activity for DHM and DHK. A study by Choudhary and Pucker [34] speculated that through a yet unknown mechanism, *DFR* with N at their 3rd position in the conserved region might become more specific for the substrate that is not favoured by the additional *DFR* copy. On the other hand, a simpler explanation is that the *DFR2* copies are undergoing neofunctionalization, evolving toward a yet unknown novel function under positive selection, while *DFR1* is maintained for its ancestral role under purifying selection. This idea is supported by the observation that the conserved region of *DFR2* copies has diverged from the ancestral *DFR*, including the insertion of a new amino acid between positions 5 and 6 (**Figure 3B**), along with several additional substitutions. However, it is still difficult to conclude that neofunctionalization is the fate in this case, as the specific new function of *DFR2* has not yet been identified. An alternative explanation is escape from adaptive conflict (EAC). In this model, the ancestral single-copy *DFR* in *Fagopyrum* may have been under selection to perform both a novel function and the ancestral *DFR* function. Because improvements to one function would negatively affect the other due to pleiotropic constraints, duplication was favoured [19]. This resulted in a *DFR2* copy free to optimize the novel function, and a *DFR1* copy maintained for the ancestral function. However, this case does not fully meet the criteria set by Des Marais and Rauscher [19], which require that both duplicates undergo adaptive changes and that the ancestral function itself is improved. Here, only *DFR2* shows adaptive changes in the conserved region, while *DFR1* appears to be maintained under purifying selection, with no evidence that the ancestral function has been enhanced.

In conclusion, the fate of the *DFR* duplicates cannot yet be determined with certainty. The most plausible hypothesis may be EAC, supported by Des Marais and Rauscher who concluded that EAC is clearly a feature of *DFR* evolution [19]. However, further experimental studies are needed to confirm whether *DFR2* has acquired a novel function and if *Fagopyrum DFR1* indeed has an improved ancestral function. Following duplication, some level of functional redundancy is expected. However, redundancy may persist not because it is beneficial, but because insufficient time has passed for its loss. Whether the duplicates like *FeDFR2b*, *FdDFR2a*, and *FdDFR2b* that show no expression, can be considered a pseudogene remains unresolved. The gene does not contain any inactivating mutations such as premature stop codons or frameshifts. Nonfunctional duplicates are not always deleted immediately as some may have only recently begun pseudogenisation and are in the early stages of decay [2]. Alternatively, the duplicates may still be retained because not enough time has passed for deleterious mutations to accumulate and eliminate it fully. Maere et al. [107] suggested that genes involved in secondary metabolism and biotic stress responses decay more slowly after both small- and large-scale duplication events. This could explain why DFR duplicates are still retained, as DFR is part of the flavonoid pathway and may provide adaptive diversity in secondary metabolites.

## List of Abbreviations

DFR: dihydroflavonol-4-reductase
DHK: dihydrokaempferol
DHQ: dihydroquercetin
DHM: dihydromyricetin
DDAI: DNA duplication associated with inversion
WGD: whole-genome duplication
TE: Transposable element
EAC: escape from adaptive conflict
TSS: transcription start site
NHEJ: non-homologous end joining

## Declaration

### Ethics approval and consent to participate

Not applicable.

### Consent for publication

Not applicable.

### Availability of data and materials

The datasets supporting the result and conclusions of this article are included in the article and additional files or are available from public databases. All the sequences and scripts used in the different analyses have been deposited in the author’s GitHub repository (https://github.com/SyafiqSamsolnizam/DFR-of-Fagopyrum).

### Competing interests

The authors declare that they have no competing interests.

### Funding

This work received no external funding.

### Authors’ contributions

SS and BP conceptualized the research project. SS carried out the analyses and prepared figures. SS wrote the manuscript. SS and BP revised the manuscript. The authors read and approved the final manuscript.

## Acknowledgements

We thank the members of the Plant Biotechnology and Bioinformatics group at the University of Bonn for their helpful discussions and valuable comments during the analysis and revision of this manuscript. This work was supported by the de.NBI Cloud within the German Network for Bioinformatics Infrastructure (de.NBI) and ELIXIR-DE (Forschungszentrum Jülich and W-de.NBI-001, W-de.NBI-004, W-de.NBI-008, W-de.NBI-010, W-de.NBI-013, W-de.NBI-014, W-de.NBI-016, W-de.NBI-022).

## Additional File

Additional file 1 (.pdf) : List of datasets used in this study to retrieve the gene of *DFR*.

Additional file 2 (.pdf) : Extended version for the amino acid and CDS multiple sequence alignment of DFR.

Additional file 3 (.pdf) : Supporting maximum likelihood tree of DFR sequences constructed using FastTree and RAxML-NG.

Additional file 4 (.xls) : SRA run IDs for gene expression analysis.

Additional file 5 (.xls) : Comparison of substitution rates between orthologous and paralogous DFR genes among *Fagopyrum* species.

Additional file 6 (.pdf): Dot plots of the genomic loci of *DFR* copies in *F. dibotrys* and *F. homotropicum*.

## Notes

https://github.com/SyafiqSamsolnizam/DFR-of-Fagopyrum

### Competing Interest Statement

The authors have declared no competing interest.

