## Additional file 1 for "Expansion of the *DFR* Gene Family in *Fagopyrum*"

Additional file 1: List of datasets used in this study to retrieve the gene of *DFR*.

| **Species** | **Data set / Genome ID** | **Database** | **Reference** |
|---|---|---|---|
| *Fagopyrum esculentum* | GCA_033239045.1 | NCBI | [1] |
| *Fagopyrum tataricum* | GWHFQEG00000000.1 | NGDC, CNCB | [2] |
| *Fagopyrum homotropicum* | GCA_033239125.1 | NCBI | [1] |
| *Fagopyrum dibotrys* | - | Figshare | [3] |
| *Persicaria maculosa* | GCA_963922045.1 | NCBI | [4] |
| *Persicaria tinctoria* | GCA_037127255.1 | NCBI | [5] |
| *Rheum nobile* | GCA_027886185.1 | NCBI | [6] |
| *Oxyria digyna* | GCA_029168935.1 | NCBI | [8] |
| *Polygonum maritimum* | GCA_963924305.1 | NCBI | [9] |
| *Polygonum aviculare* | GCA_934048045.1 | NCBI | [10] |
| *Fallopia multiflora* | GCA_041753945.1 | NCBI | [11] |
| *Reynoutria japonica* | GCA_048128825.1 | NCBI | [12] |
| *Rheum palmatum* | - | Figshare | [13] |
| *Rheum tanguticum* | - | Figshare | [14] |
| *Rumex hastatulus* | 65183 | CoGe | [15] |
| *Rumex scutatus* | GCA_046244935.1 | NCBI | [16] |
| *Rumex bucephalophorus* | GCA_049309485.1 | NCBI | [17] |
| *Fallopia convolvulus* | GCA_964016875.1 | NCBI | [18] |
