## Additional file 2 for "Expansion of the *DFR* Gene Family in *Fagopyrum*"

Additional file 2: (A) Extended version for multiple sequence alignment including *FeDFR1a* & *FeDFR2* from the study of Katsu et al. (B) Multiple sequence alignment of the CDS in the conserved region of each DFR duplicates.

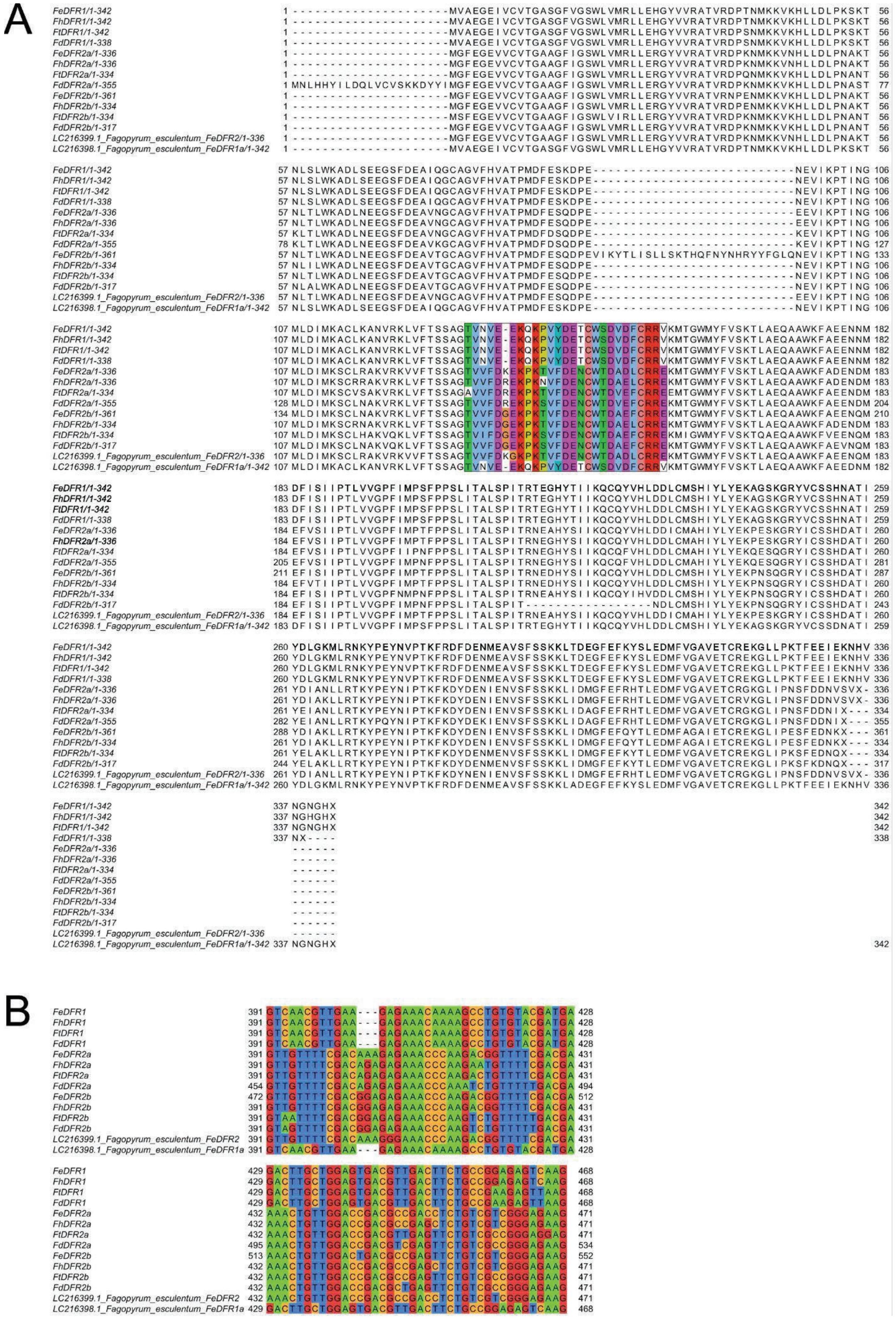
