## Additional file 3 for "Expansion of the *DFR* Gene Family in *Fagopyrum*"

Additional file 3: Maximum likelihood tree of DFR sequences constructed using (A) FastTree with standard parameters and (B) RAxML-NG with the model TIM3+G4, 1000 bootstrap replicates, and 50 starting trees.

A

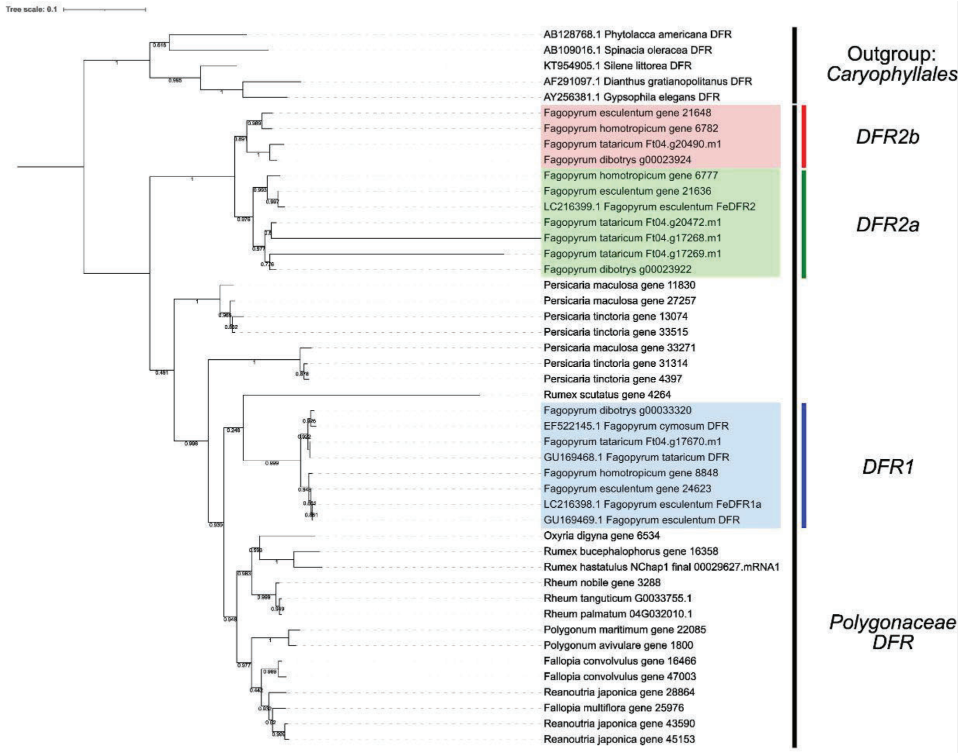


B

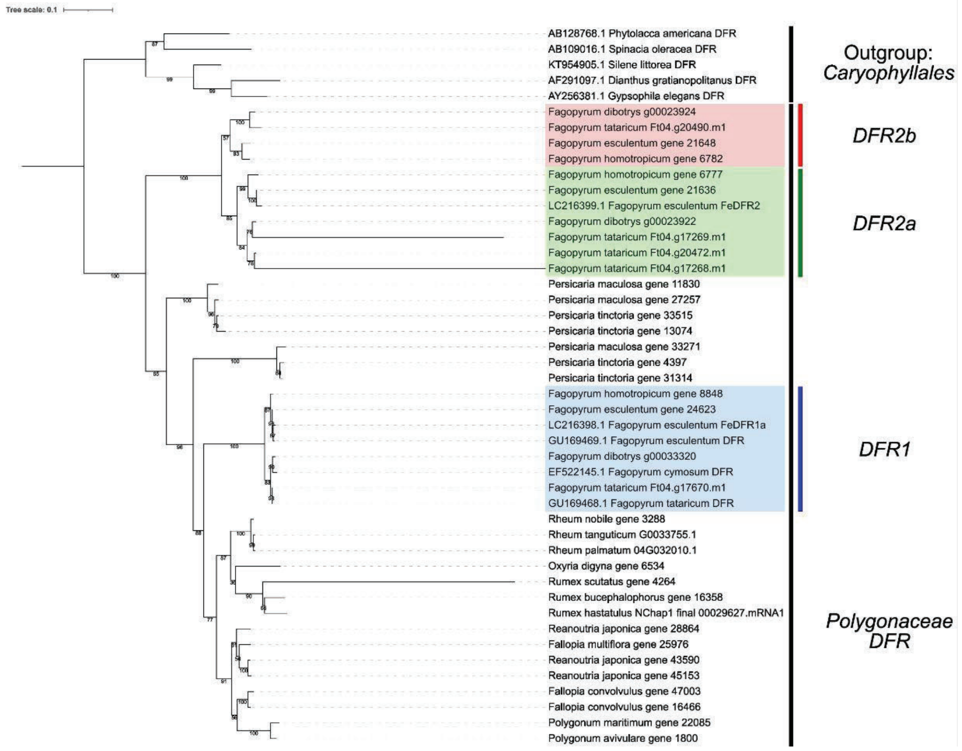
