## Additional file 6 for "Expansion of the *DFR* Gene Family in *Fagopyrum*"

Additional file 6: Dot plots of the genomic loci of DFR copies in *F. dibotrys* and *F. homotropicum*. Each panel represents a pairwise alignment of genomic regions containing DFR genes. The chromosome identifiers and location are shown along the axes. The gene annotation is included outside of the axes, with vertical and horizontal lines marking the positions of DFR genes. A & B = *DFR1* loci, C & D = *DFR2* loci, E & F = *DFR1* & *DFR2* loci.

*F. dibotrys* *F. homotropicum*

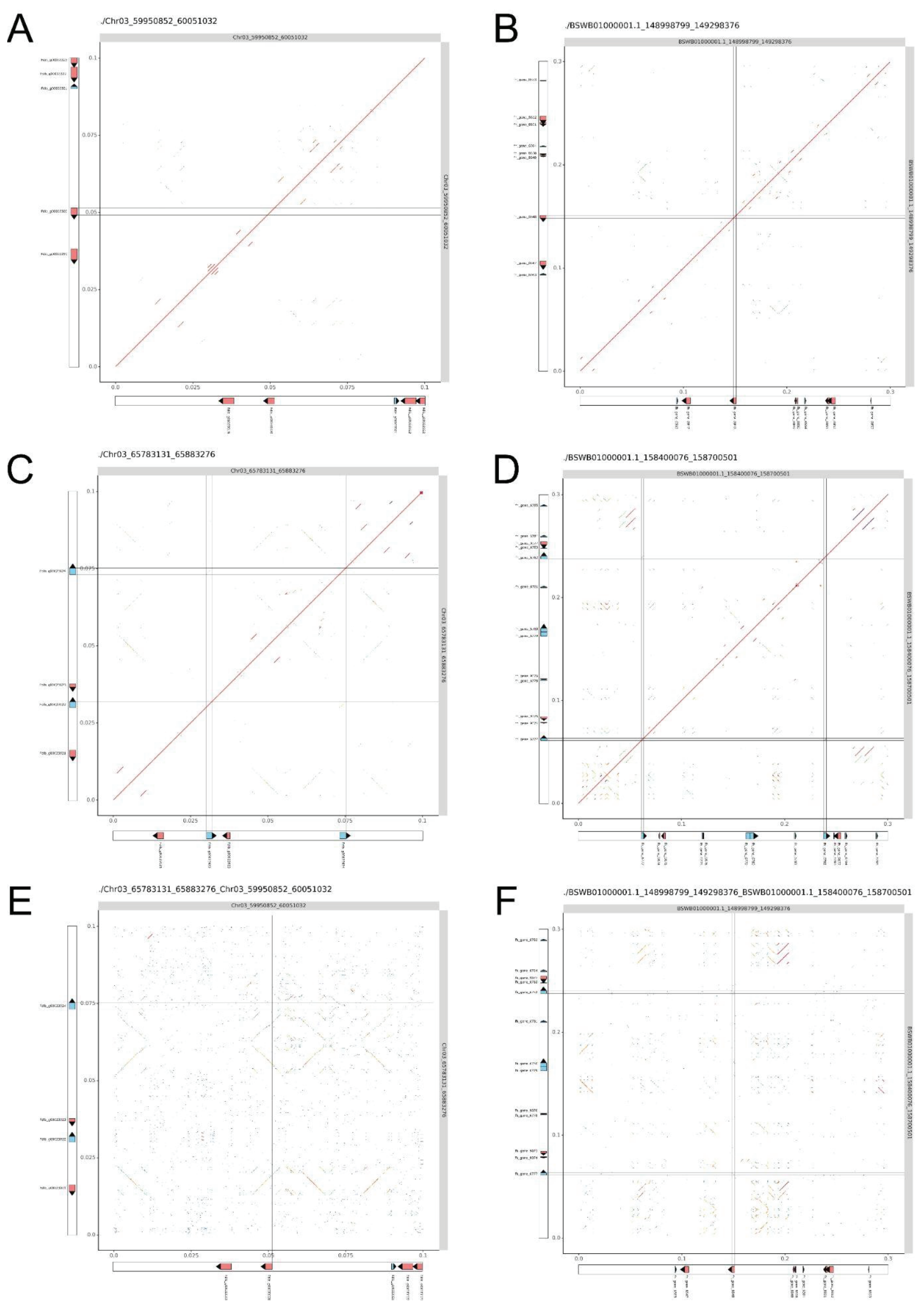
